# Single-particle tracking reveals heterogeneous PIEZO1 diffusion

**DOI:** 10.1101/2022.09.30.510193

**Authors:** Alan T. Ly, J. Alfredo Freites, Gabriella A. Bertaccini, Elizabeth L. Evans, George D. Dickinson, Douglas J. Tobias, Medha M. Pathak

## Abstract

The mechanically-activated ion channel PIEZO1 is critical to numerous physiological processes, and is activated by diverse mechanical cues. The channel is gated by membrane tension and has been found to be mobile in the plasma membrane. We employed single particle tracking (SPT) of endogenous, tdTomato-tagged PIEZO1 using Total Internal Reflection Fluorescence Microscopy in live cells. Application of SPT unveiled a surprising heterogeneity of diffusing PIEZO1 subpopulations, which we labeled “mobile” and “immobile”. We sorted these trajectories into the two aforementioned categories using trajectory spread. To evaluate the effects of the plasma membrane composition on PIEZO1 diffusion, we manipulated membrane composition by depleting or supplementing cholesterol, or by adding margaric acid to stiffen the membrane. To examine effects of channel activation on PIEZO1 mobility, we treated cells with Yoda1, a PIEZO1 agonist, and GsMTx-4, a channel inhibitor. We collected thousands of trajectories for each condition, and found that cholesterol removal and Yoda1 incubation increased the channel’s propensity for mobility. Conversely, we found that GsMTx-4 incubation and cholesterol supplementation resulted in a lower chance of mobile trajectories, whereas margaric acid incubation did not have a significant effect on PIEZO1 mobility. The “mobile” trajectories were analyzed further by fitting the time-averaged mean-squared displacement as a function of lag time to a power-law model, revealing mobile PIEZO1 puncta exhibit anomalous subdiffusion. These studies illuminate the fundamental properties governing PIEZO1 diffusion in the plasma membrane and set the stage to determine how cellular processes and interactions may influence channel activity and mobility.

**SIGNIFICANCE:** PIEZO1 is a mechanically-activated ion channel that regulates a number of physiological processes. Here we examine a fundamental biophysical property of PIEZO1 - its movement in the plasma membrane. We find that the mobility of PIEZO1 is surprisingly heterogeneous, with some PIEZO1 puncta showing high mobility and some displaying very limited mobility. Cholesterol depletion from the plasma membrane increases PIEZO1 mobility while cholesterol supplementation decreases mobility. Yoda1 treatment increases PIEZO1 mobility whereas GsMTx-4 treatment decreases channel mobility.

## INTRODUCTION

Mechanically-activated ion channels rapidly sense and transduce mechanical stimuli into electrical and chemical signals by allowing ion flux across biological membranes, and are found across bacteria, archaea, and eukaryotes (1–5). The PIEZO family of ion channels was the first mammalian excitatory mechanically-activated group of ion channels to be identified (6). The importance of PIEZO channels is underscored by the fact that they are highly conserved across many species, and are expressed in a wide range of tissues. PIEZO channels activate in response to mechanical cues and cause cationic influx (6), thereby regulating a number of crucial biological processes. These proteins are critical in vascular development (7, 8), lymphatic valve development (9), bone formation (10), blood pressure baroreflex (11), mechanical itch (12) and touch (13, 14), proprioception (15), tactile and mechanical pain (16, 17), skin wound healing (18), and neural stem cell differentiation (19). Knockouts of PIEZO1 are embryonic lethal (7, 8), and PIEZO1 mutations are associated with several diseases, including dehydrated hereditary stomatocytosis, and lymphatic dysplasia (20, 21).

The homotrimeric PIEZO channels, with their triple-bladed, propeller-like architecture, has a unique structure compared to other known membrane proteins (22–26). The propeller blades consist of repeating four-transmembrane-helix-containing bundles that are linked to the central pore by the beam and anchor domains (22). Structural, computational, and microscopy studies of PIEZO channels reveal that the channel structure causes local distortion of the membrane, thereby inducing membrane curvature and causing the membrane to adopt a striking bowl-like characteristic (24, 26–30). Membrane tension gates PIEZO1 (31–33), demonstrating that the channel directly senses forces on the lipid bilayer. PIEZO1 has also been proposed to function through a force-through-filament mechanism, in which the actin cytoskeleton may function as a tether to translate mechanical forces from the membrane to PIEZO1 to induce conformational change, thereby gating the channel (34, 35).

We previously reported that cellular traction forces generated by the actomyosin cytoskeleton and transmitted to the substrate at focal adhesions can activate PIEZO1 (19, 36). Using a PIEZO1-tdTomato reporter mouse model where the endogenous PIEZO1 protein is tagged with a tdTomato fluorophore at the PIEZO1 C-terminus, we found that the PIEZO1 protein localization is not restricted to focal adhesions and that the channel is surprisingly mobile within the plasma membrane (36). Ridone *et al*. similarly found that the channel was mobile using heterologously-expressed PIEZO1-GFP, and further showed that cholesterol depletion via methyl-*β*-cyclodextrin (MBCD) increased channel diffusion and disrupted clustering of PIEZO1 (37). Earlier analysis was performed under the assumption that PIEZO1 demonstrated Brownian motion (37). However, the plasma membrane through which PIEZO1 diffuses is a complex environment composed of a number of proteins and lipids with considerable structural heterogeneity, which could influence PIEZO1 mobility (38–40). Indeed, in a later study Vaisey *et al*. observed that PIEZO1 in red blood cells demonstrated a confined Brownian motion (41).

Here, we report single-particle tracking (SPT) of endogenously expressed PIEZO1-tdTomato channels. Visual examination reveals heterogeneous trajectories that could be classified into two broad categories based on their spatial extent: “mobile” class wherein trajectories displayed relatively large spatial extent, and “immobile” trajectories limited to a small area. We show that PIEZO1-tdTomato is more likely to be classified as “mobile” when the cells are treated with Yoda1 and MBCD. Conversely, the channel is more likely to be “immobile” when treated with GsMTx-4 or when the membrane is supplemented with cholesterol. The “mobile” class was also found to be subdiffusive across all the tested experimental conditions. Our results demonstrate that membrane composition and channel activity may play a key role in regulating PIEZO1 mobility.

## MATERIALS AND METHODS

### ANIMALS

The University of California, Irvine and its associated Institutional Animal Care and Use Committee has approved all studies performed in this study. All experiments were performed in accordance to the guidelines. All cells used in this study were harvested from a reporter mouse with a tdTomato knock-in at the C-terminus of the endogenous PIEZO1 channel (JAX stock 29214) (7).

### Mouse Embryonic Fibroblast (MEF) Isolation and Culture

MEF cells were isolated from the tdTomato knock-in reporter mice (7). Mice were considered embryonic day 0.5 upon vaginal plugging. Fibroblast cells were harvested from embryos at embryonic day 12.5 after removing the head, limbs, and tail from the embryo. The dissection was performed in 33 mM D-(+)-glucose (Sigma-Aldrich, G-6152) and 1% Penicillin-Streptomycin (10,000 U/mL; Gibco, 15140122) in Dulbecco’s Phosphate-Buffered Saline (Gibco, 14-190-250). The brain was harvested for mouse Neural Stem Cells (mNSC) (see **Mouse Neural Stem Cell (mNSC) Isolation and Culture** below). The tissue was spun at 260g for 5 min, and the supernatant was aspirated. The cells were resuspended in DMEM (ThermoFisher Scientific, 11960-051) with 15% fetal bovine serum (Omega Scientific, FB-12), 1x GlutaMax (ThermoFisher Scientific, 35050-061), 1 mM sodium pyruvate (ThermoFisher Scientific, 11360-070), and 1x non-essential amino acid solution (ThermoFisher Scientific, 11140-050). Cells were plated in a T-25 cell culture flask (Eppendorf, 0030710126) coated with 0.1% gelatin solution (Fisher Scientific, ES-006-B) and incubated in a sterile environment at 37°C with 5% CO_2_. Media was changed 1 hour after plating. PIEZO1-tdTomato MEFs were passaged using TrypLE Express (ThermoFisher, 12604013) to dissociate the cells and were spun at 260g for 5 min. Cells were then counted using a hemocytometer and 7,500-10,000 cells were plated on the 14mm glass region of #1.5 glass-bottom dishes (Mat-Tek Corporation) coated with 10 µg/mL fibronectin (Fisher Scientific, CB-40008A). Media was changed after 2h and every 48h until imaging experiments. Cells were maintained in a 5% CO_2_ incubator at 37ºC for at least 72h prior to imaging. MEFs were used for experiments between passages 3 and 7.

### Mouse Liver Sinusoidal Endothelial Cell (mLSEC) Isolation and Culture

mLSECs were isolated from 8-week old PIEZO1-tdTomato reporter mice using an immunomagnetic separation technique. A mouse liver was thoroughly minced using scalpel blades and resuspended in a dissociation solution containing 9 mL 0.1% collagenase II, 1 mL 2.5 U ml-1 dispase, 1 µM CaCl_2_ and 1 µM MgCl_2_ in Hanks Buffer solution. The tissue-dissociation mix was incubated at 37ºC for 50 mins in a tube rotator to provide continuous agitation. Following this enzymatic digestion, the mix was passed through 70 and 40 µm cell strainers to remove undigested tissue. Cells were washed twice in PEB buffer containing phosphate-buffered saline solution (PBS), EDTA 2mM and 0.5% BSA, pH 7.2. The washed pellets were resuspended in 1 mL PEB buffer and 30 µL CD146 microbeads (Miltenyi Biotech) at 4ºC for 15 min under continuous agitation. CD146 is a membrane protein marker for endothelial cells and is highly expressed in mLSECs. Following incubation, the solution was passed through an LS column (Miltenyi Biotech) primed with PEB buffer. The column was washed 3 times with 5 mL PEB buffer and the CD146 negative eluate was removed. CD146 positive cells were retained in the column and flushed with 5 mL warmed EGM-2 growth medium supplemented with EGM-2 bullet kit (Lonza) into a separate tube. Cells were spun at 300g for 5 min, diluted in 1 mL EGM-2 media and counted using a hemocytometer. 30,000-40,000 cells were plated on the 14 mm glass region of #1.5 glass-bottom dishes (Mat-Tek Corporation) coated with 10 µg/mL fibronectin (Fisher Scientific, CB-40008A). Media was changed after 2h and every 48h until imaging experiments. Cells were grown in a 5% CO_2_ incubator at 37ºC for at least 72h prior to imaging.

### Mouse Neural Stem Cell (mNSC) Isolation and Culture

mNSCs were isolated from the PIEZO1-tdTomato knock-in reporter mouse (7). Embryos were obtained on embryonic day 12.5, and heads of the mice were harvested in 33 mM D-(+)-glucose (Sigma-Aldrich, G-6152) and 1% Penicillin-Streptomycin (10,000 U/mL; Gibco, 15140122) in Dulbecco’s Phosphate-Buffered Saline (Gibco, 14-190-250). The top layer of the head was removed to visualize the cortex, and the top half of each cortex was harvested and placed on ice. Tissue was spun at 260g for 5 min, the supernatant was discarded. Cells were resuspended in Dulbecco’s modified Eagle’s media (Thermo Fisher Scientific, 11995-065), 1× N2 (Thermo Fisher Scientific, 17502048), 1× B27 (Thermo Fisher Scientific, 17504044), 1 mM sodium pyruvate (Thermo Fisher Scientific, 11360070), 2 mM Glutamax (Thermo Fisher Scientific, 35050061), 1 mM N-acetylcysteine (Millipore Sigma, A7250), 10 ng/ml b-FGF (Peprotech, 100-18B), 20 ng/ml EGF (Peprotech, AF-100-15), and 2 µg/ml heparin (Millipore Sigma, H3149). mNSCs were cultured as neurospheres in non-adherent cultureware, and were passaged using Neurocult Chemical Dissociation Kit (Stem Cell Technologies, 05707). 10,000 mNSCs were plated onto the 14mm glass region of #1.5 glass-bottom dishes (Mat-Tek Corporation) coated with 10 µg/mL laminin (Thermo Fisher Scientific, 23017015). mNSCs between passages 4-7 and days 29-33 were used for imaging. mNSCs were cultured in a 5% CO_2_ incubator at 37ºC for at least 72h prior to imaging.

### Imaging PIEZO1-tdTomato

Mobility of native PIEZO1-tdTomato channels was imaged using Total Internal Reflection Fluorescence (TIRF) microscopy at 37°C. PIEZO1-tdTomato MEFs, mLSECs, and mNSCs were washed with phenol red-free DMEM/F12 (Invitrogen, 25116001) thrice and incubated in imaging solution, composed of 148 mM NaCl, 3 mM CaCl_2_, 1 mM KCl, 2 mM MgCl_2_, 8 mM Glucose, 10 mM HEPES, pH 7.30, and 316 mOsm/L osmolarity for 5 min. An Olympus IX83 microscope fitted with a 4-line cellTIRF illuminator, an environmental control enclosure and stage top incubator (Tokai Hit), and a PLAPO 60x oil immersion objective NA 1.45 were used to image cells. A programmable motorized stage (ASI) was used to identify samples throughout imaging. Images were acquired using the open source software µ-Manager (42). Cells were illuminated with a 561 nm laser and images were acquired using a Hamamatsu Flash 4.0 v2+ scientific CMOS camera at a frame rate of 10 frames/second with a 100 ms exposure time.

PIEZO1-tdTomato MEFs were fixed using a 4% paraformaldehyde (Electron Microscopy Sciences, 15710), 1x PBS, 5 mM MgCl_2_, 10 mM EGTA, 40 mg/mL sucrose buffer for 10 min at room temperature. The cells were washed thrice with PBS for 5 minutes.

### Drug Treatment

Methyl-*β*-cyclodextrin-treated cells were incubated in 10 mM methyl-*β*-cyclodextrin (Sigma-Aldrich, C4555-5G) for 15 min before imaging. Cholesterol-MBCD-treated cells were incubated in 100 µg/mL Cholesterol-water soluble (containing methyl-*β*-cyclodextrin for solubility) (Sigma-Aldrich, C4951-30mg) and were incubated for 1 hour. Cells treated with 150 µM margaric acid (NuChek N-17-A) were incubated overnight at 37°C for 24h. Cells treated with 4 µM Yoda1 (Tocris 558610) or 4 µM GsMTx-4 (Tocris 4912) were incubated for 15 min. Margaric acid and Yoda1 were dissolved in Dimethyl sulfoxide (DMSO, Sigma Aldrich 276855-100ML). As such, DMSO was used as a control for margaric acid and Yoda1. Imaging sessions were limited to 30 min after drug treatment, and the chemical agents were maintained in the bath solution during imaging.

### PIEZO1-tdTomato Trajectory Generation

Single particle tracking of PIEZO1-tdTomato puncta was done using the custom-built, open-source image processing and analysis program Flika (43) on Python 3.9.13. A difference of Gaussians algorithm was used as a spatial bandpass filter on the image stacks. The resulting enhanced stack was then thresholded using a manually selected threshold value to generate a binary image stack. Spatially-continuous pixels above this threshold were considered a single particle. A two-dimensional Gaussian function was used to determine the centroid of each particle to subpixel precision. Particles within three pixels of consecutive frames were assumed to represent the same PIEZO1-tdTomato puncta. These particles were then linked to generate trajectories. Average nearest neighbor per frame and step sizes were calculated to verify that ID switching does not occur for the majority of trajectory linkages (See Supplemental Methods for details). Skipped frames were handled by inserting a placeholder value (numpy.nan) for missing coordinates (36, 44). A conversion factor equivalent to the length of a single pixel, 0.1092 µm, was used to transform two-dimensional coordinates in pixel units to microns. We limited our analysis to trajectories that were at least 20 seconds in length, which at a frame rate of 100 milliseconds, resulted in a minimum of 200 positions per trajectory. Trajectory analyses were performed with R (version 4) (45), unless stated otherwise.

### Fixed-cell trajectory analysis

We used fixed-cell trajectories to estimate the magnitude of the localization error under the assumption that their apparent spread should stem, exclusively, from the localization error (46). To identify trajectories consistent with this assumption, we first computed the individual trajectory time averaged mean square displacement (TAMSD), 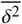, as a function of time of according to 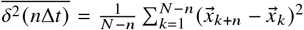 for an *N*-point trajectory, 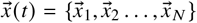, sampled at a frame rate Δ*t* The TAMSD computations were performed over one decade of frame lags *n* = (1, …, 10), and their time dependence modeled as a power-law, 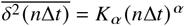, over the second to fourth frame lags (*n* = 2, 3, 4), using linear regression on the log-transformed variables. Under our assumption, we expect *α* ≈ 0. We identify trajectories that fulfill this criterion by imposing a two-component Gaussian mixture model over the resulting power-law exponent, *α*, distribution and choosing the sub-population with lower mean value of *α*. We refined our data set further by imposing a 2-component kernel-based mixture model over the resulting distribution of power-law coefficients, *K*_*α*_, and choosing the sub-population with lower mean value of *K*_*α*_. Both two-component mixture models were generated using expectation minimization algorithms implemented in the mixtools R package (47). A joint (*K*_*α*_, *α*) kernel density estimate of the resulting trajectory data set indicated that most of the sampled trajectories were consistent with our initial assumption. Our estimate for the magnitude of the localization error was taken as the most likely *K*_*α*_ value according to the joint kernel density estimate, which under the assumption *α* ≈ 0 is 0.33 Å^2^.

We also computed the corresponding scaled radius of gyration, sR_g_, (see Eq. 1) for this selected data set of fixed-cell trajectories and used the 95^th^ percentile of corresponding distribution as a threshold to identify trajectories as immobile or mobile in live cells.

### Live-cell trajectory analysis

Prior to performing the analyses reported here, we computed the individual trajectory TAMSD up to 20 frame lags (see previous section) and used the results to remove any trajectory with TAMSD values below our localization error estimate.

An *N*-point trajectory with position vectors 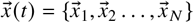 can be equivalently described in terms of step vectors, 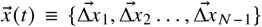, with 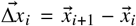. We used this step vector representation to characterize the live-cell trajectories mobile class. To probe heterogeneity at the mobile class ensemble level, we modeled the entire one-dimensional steps distribution as a single Gaussian, using the sample mean and variance; and as a two-component Gaussian distribution, using Mathematica 14 (48) to obtain the maximum likelihood parameter estimates. To probe heterogeneity at the individual mobile trajectory level, we modeled the distribution of step vector magnitudes as a mixture distribution with one through four Rayleigh distribution components. We used Mathematica (48) to obtain the maximum likelihood parameter estimates for each candidate mixture distribution. To select the number of components in the mixture distribution of each individual trajectory, we used Akaike information criterion statistic (AIC), given by 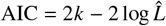 with *K* the number of degrees of freedom and 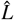is the maximum likelihood estimate, and selected either the model with the minimum AIC value or, if the model with minimum AIC had (*j* + 1) -components and AIC_*j*_ − AIC_*j*+1_ < 4, we chose the model with *j*-components.

To assess the effect of different drug treatments on the mobility of PIEZO1, we used the number of mobile and immobile trajectories in a given experimental session to compute an estimate of the odds of observing mobile trajectories and used the results to obtain odds ratio estimates of observing mobile trajectories upon drug treatment relative to their corresponding control. For drug treatments in aqueous solution, we grouped drug-treated and control trajectories from the same experimental session and used a Mantel-Haenszel significant test to compute a common odds ratio estimate using the total number of drug-treated and control trajectories in each experimental session as weights. For drug treatments in DMSO, we lacked control trajectories from the same experimental session as the drug treatments. Therefore, we constructed the common odds ratio estimate using the total number of trajectories in the drug-treated groups as weights.

The time dependence of individual trajectories’ TAMSD was modeled as a power-law over the first 20 frame lags using linear regression on the log-transformed independent and dependent variables. Individual trajectories’ TAMSD and the localization error estimate from the fixed-cell data were used to obtain ensemble estimates of the power-law exponent using the procedure for noisy and heterogeneous trajectories described in ref. (49).

The code used for the trajectory analysis is available in an online repository (see Supplemental Material).

## RESULTS

### PIEZO1-tdTomato puncta exhibit heterogeneous mobility

We imaged with TIRFM endogenously expressed PIEZO1-tdTomato in mouse embryonic fibroblast cells (MEFs) harvested from PIEZO1-tdTomato reporter mice. PIEZO1 channels were visible as distinct puncta as described earlier (36) but with higher fidelity due to improvements in camera technology (Fig. 1A). Ellefsen *et al*. had previously acquired PIEZO1 diffusion videos using an Andor iXon3 electron multiplying charge-coupled device (EMCCD) camera. We captured our videos using a Hamamatsu Flash 4.0 v2+ scientific Complementary Metal Oxide Semiconductor (sCMOS) camera, which have several advantages over EMCCDs in pixel size/resolution, signal:noise, speed, dynamic range, and a larger field of view (50–52). Visual inspection of videos collected revealed that some puncta were quite mobile while others showed little or no mobility (Fig. 1A and Supplemental Video 1 in Supplemental Material). The reduced mobility of some PIEZO1 puncta was particularly evident in regions of the cell where PIEZO1 puncta appeared to cluster together in structures reminiscent of focal adhesions (Fig. 1A, green inset; also compare green and blue insets in Supplemental Video 1), in agreement with reports that PIEZO1 is enriched at focal adhesions under certain conditions (53, 54). We examined individual PIEZO1 trajectories more closely and found that they could be classified according to their apparent two-dimensional extent into a “mobile” class and “immobile” class (Fig.1B, 1C).

**Figure 1:**
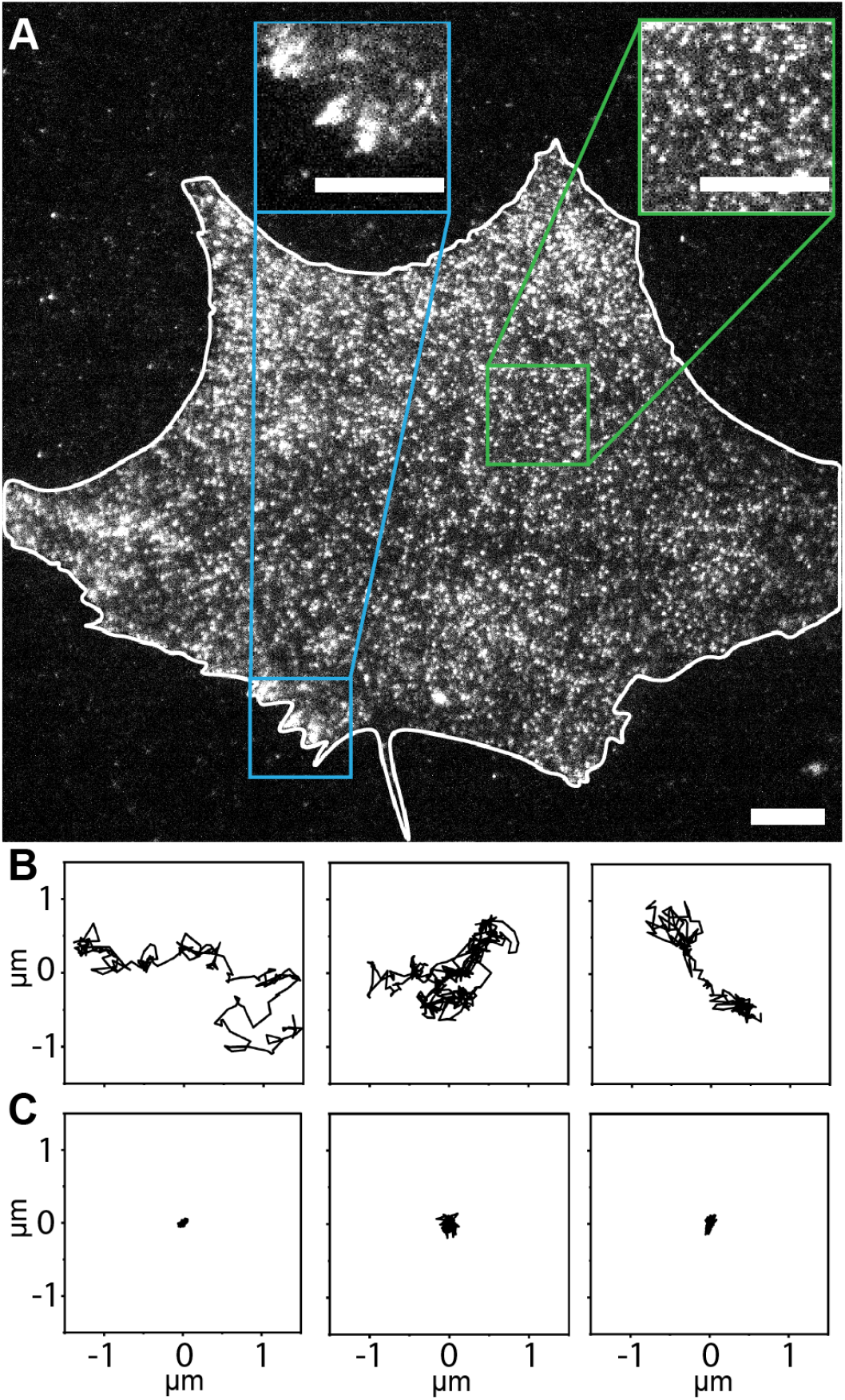
TIRF microscopy and single particle tracking reveals heterogeneity in PIEZO1-tdTomato mobility. A. Representative TIRF image of PIEZO1-tdTomato puncta in live MEFs harvested from PIEZO1-tdTomato reporter mice. The white line denotes the cell boundary. Insets show enlarged regions of interest. The green inset is representative of regions where puncta appear mobile, whereas the blue inset is representative of regions where puncta show little or no mobility. The trajectories generated from the single particle tracking analysis can be classified into a (B) mobile fraction, and (C) an immobile fraction, according to their spatial spread (see Fig. 2). See also Supplemental Video 1 in Supplementary Material. Scale bars = 10 µm.

To classify trajectories as “mobile” or “immobile”, we made the assumption that the apparent spread of immobile trajectories should stem exclusively from the localization error. We extracted trajectories from paraformaldehyde-treated, fixed cells, in which PIEZO1-tdTomato puncta are rendered immobile (See Supplemental Video 2). To quantify the apparent spread of a two-dimensional trajectory, we used the trajectory’s radius of gyration scaled by the corresponding mean step-length, as proposed by Golan and Sherman (46):

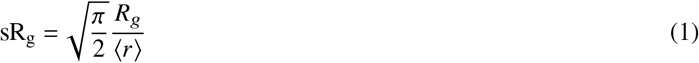

where for an *N*-point trajectory 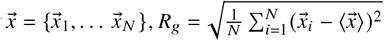 with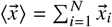 and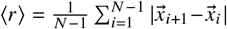. As shown in ref. (46), positions sampled from a two-dimensional isotropic Gaussian lead to 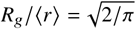. Thus, for immobile trajectories with apparent motion resulting exclusively from localization error, the sR_g_ distribution should approximate a Gaussian distribution with mean of 1 independent of the magnitude of the localization error (see Fig. 2A). Importantly, this relationship holds true regardless of the magnitude of localization error and thus can be used to identify immobile particles. Similar to Golan and Sherman (46), we used the 95^th^ percentile of a selected set of fixed-cell trajectories sR_g_ distribution as a threshold to categorize trajectories as immobile or mobile in live cells. Using this threshold criterion, 40% of the PIEZO1-tdTomato trajectories from live MEFs are identified as immobile (Fig. 2A). Notably, applying the same criterion to PIEZO1-tdTomato trajectories obtained from a different cell type, mouse liver sinusoidal endothelial cells (mLSECs), results in a similar partition between mobile and immobile trajectories, suggesting that the observed heterogeneity is not limited to PIEZO1-tdTomato expressed in MEFs (Fig. 2A). We also obtained trajectories from mouse neural stem cells (mNSCs), but due to the low expression of PIEZO1-tdTomato in these cells and the rapid photobleaching, we were only able extract a very small number. When we shortened the trajectory cutoff length from 20 s to 10 s for mNSCs, we obtained 2422 trajectories from 147 videos compared to 24152 MEF trajectories from 137 videos and 71218 mLSEC trajectories from 346 videos using 20 s cutoffs. While the number of trajectories obtained from mNSCs were an order of magnitude below MEF cells, the mNSCs also exhibited immobile and mobile subpopulations (57% mobile:43% immobile), but given the small number of trajectories, we focused on MEFs and mLSECs for further analysis.

**Figure 2:**
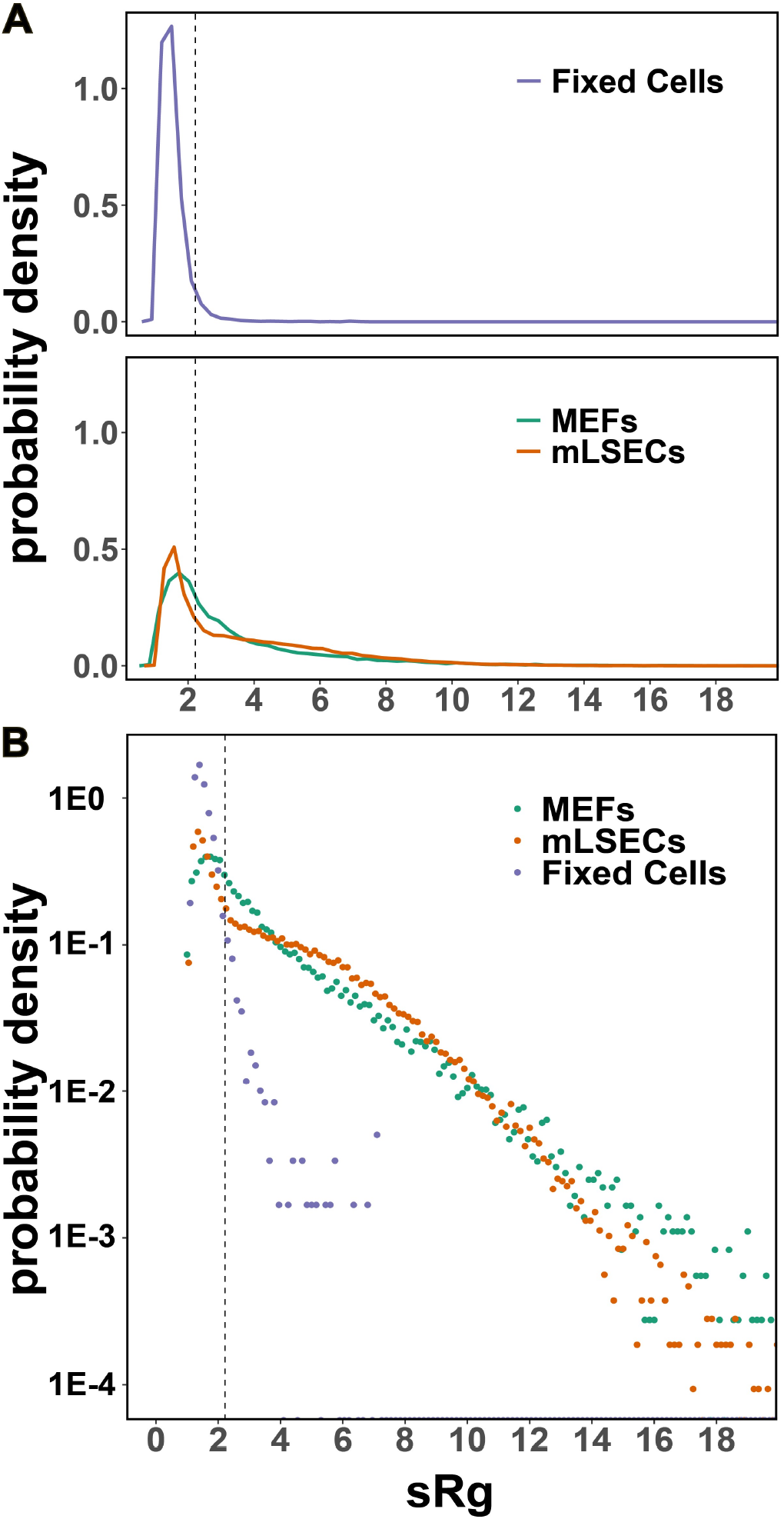
PIEZO1-tdTomato trajectories exhibit a broad range of mobilities. (A) Distribution of individual PIEZO1-tdTomato trajectories’ radius of gyration scaled by the corresponding mean step length (sR_g_) from MEFs (green), mouse liver sinusoidal endothelial cells (mLSECs, red), and MEFs fixed with paraformaldehyde. Assuming that the measured trajectory spread in fixed cells arises exclusively from the localization error, we take the 95-percentile of the MEFs fixed cells’ sR_g_ distribution (vertical dashed line) as cutoff for the separation of trajectories in live cells between a mobile class (see Fig. 1B) and an immobile class (see Fig. 1C). Using this criterion, 43% of the observed trajectories are identified as immobile in both cell types. The plot traces are joined histogram bin heights. (B) The sR_g_ distributions from live cells exhibit exponential tails for the mobile class

Next, we focused on understanding the behavior of the mobile trajectories. Unlike the distributions of sR_g_from fixed cells, the tails of the sR_g_ distributions from live cell trajectories have an exponential character (see Fig. 2B), suggesting heterogeneity within the PIEZO1-tdTomato mobile class. We confirmed this finding by considering the trajectories’ steps distribution.

Under the assumption of two-dimensional Brownian motion, the trajectory steps, (e.g. 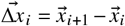 for all *i* = 1, …, *N*− 1 in an *N*-point trajectory) are distributed according to

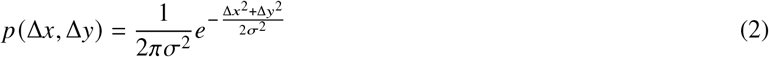

where the pair (Δ*x*, Δ*y*) denotes the components of the step vector. If a trajectory is sampled at equal time intervals, Δ*t*, the variance, *σ*^2^, can be expressed as an apparent diffusion coefficient, according to *σ*^2^ = 2*D*Δ*t*. Thus, the ensemble-level heterogeneity observed in Fig. 2B could just reflect the intrinsic heterogeneity of individual diffusers, expressed as different values of *D* for different trajectories. According to Equation (2), ⟨Δ*x*⟩^2^ = 2*D*Δ*t*, and similarly for Δ*y*. Thus, the dependence of the step distribution on individual values of *D* can be removed by scaling individual step components by the corresponding root mean square value as

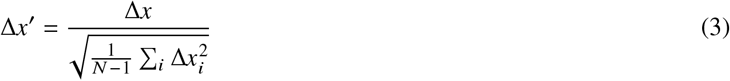

and similarly for Δ*y*(55, 56).

As shown in Fig. 3, the mobile trajectories’ step distributions in both cell types deviate from single Gaussians, and are better described by a mixture of two Gaussians, indicating that individual PIEZO1-tdTomato mobile trajectories are heterogeneous. The better fit for mLSECs is likely due to increased sampling relative to MEFs at large step lengths. This finding stands in contrast to the underlying assumption made so far in the literature that PIEZO1 diffusion can be described as Brownian motion (37, 41).

**Figure 3:**
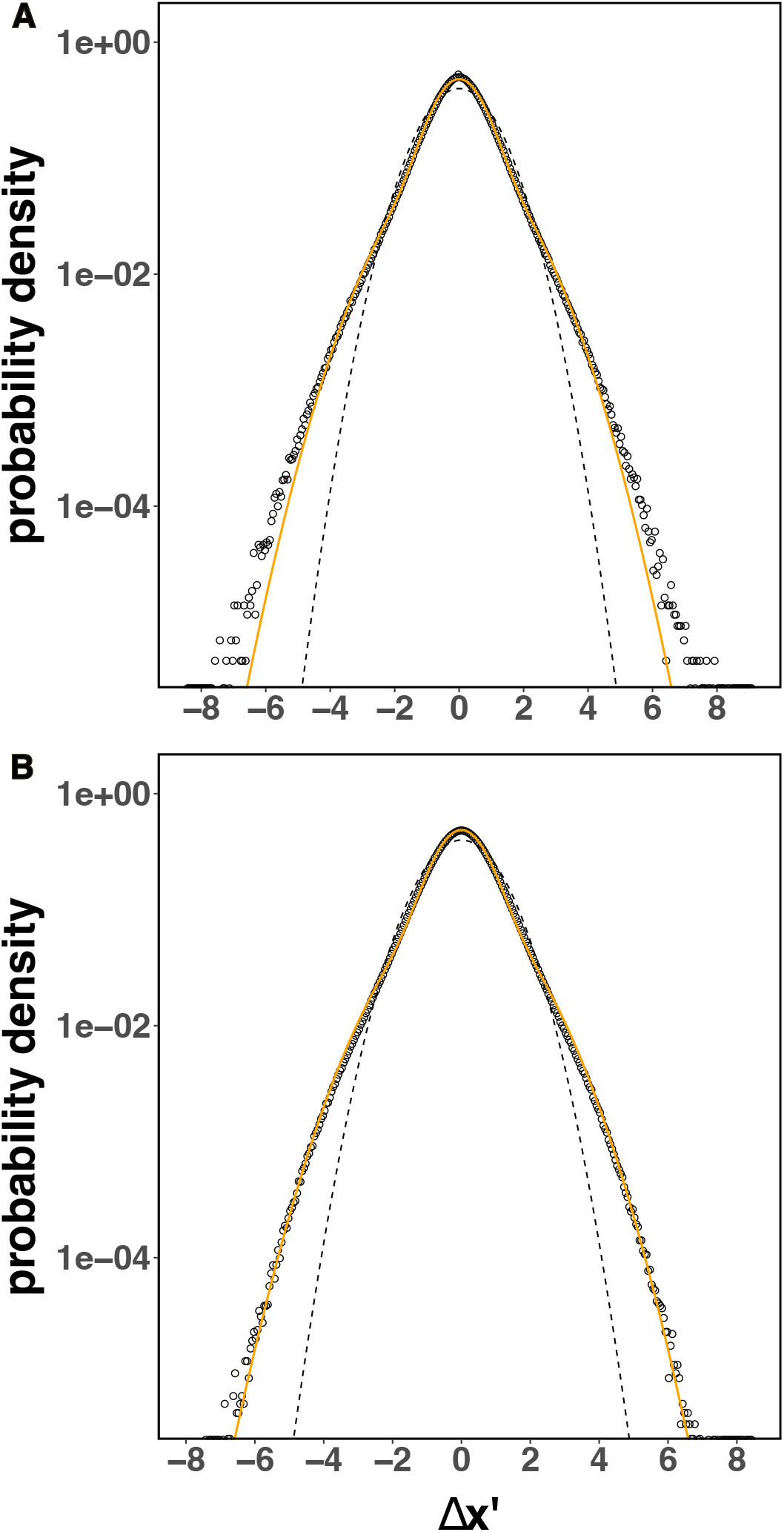
PIEZO1-tdTomato mobile trajectories exhibit non-Brownian motion. The step distributions from individual trajectories in both (A) MEFs and (B) mLSECs mobile trajectories are shown as dots (corresponding to histogram bin heights). The non-Gaussian character is evidenced when compared to a two-Gaussian mixture model (shown as continuous orange lines) and the corresponding single Gaussian distribution with same mean and variance (shown as dashed lines). The individual step vector components of each mobile trajectory, scaled by their corresponding root mean square value (see eq. 3), were binned together.

To characterize heterogeneity within the PIEZO1-tdTomato mobile class at the level of individual trajectories, we opted for a simplified description in terms of the two-dimensional step-length 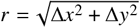. When both Δ*x* and Δ*y* are normally distributed with variance *σ*^2^, then the corresponding step-length probability density function at constant sampling rate Δ*t* is given by the Rayleigh distribution 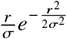. We then allow for trajectory heterogeneity by considering a simple phenomenological model for the distribution of step lengths, *p* (*r*), according to:

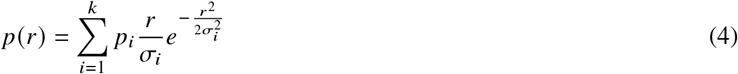

under this mixture model, an individual trajectory arises from random sampling over a finite number of mobility states *i* = 1 … *k*, implying 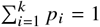 and we assume that our choice of limiting the analysis to trajectories with at least 200 positions (see Materials and Methods) provides sufficient sampling to describe the step population of each *i*^*th*^-state by a Gaussian distribution with variance 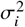. We emphasize that, as the model is intended to be strictly phenomenological, we make no assumptions on the underlying nature of these mobility states, and for a given trajectory, we take the mixing proportions, *p*_*i*_, to be stationary.

We considered models with one through four components and used maximum likelihood estimation to determine the corresponding parameters for each trajectory. An assessment of the resulting models’ Akaike information criterion statistics (57) (see Materials and Methods) indicated that the step-length distribution of most trajectories (∼ 90%) in the mobile class from each cell type could be adequately described by a two-component mixture model. Despite this uniformity in the number of components, we find that both the component proportion and apparent diffusion coefficient values are broadly distributed across trajectories, as shown in Fig. 4.

**Figure 4:**
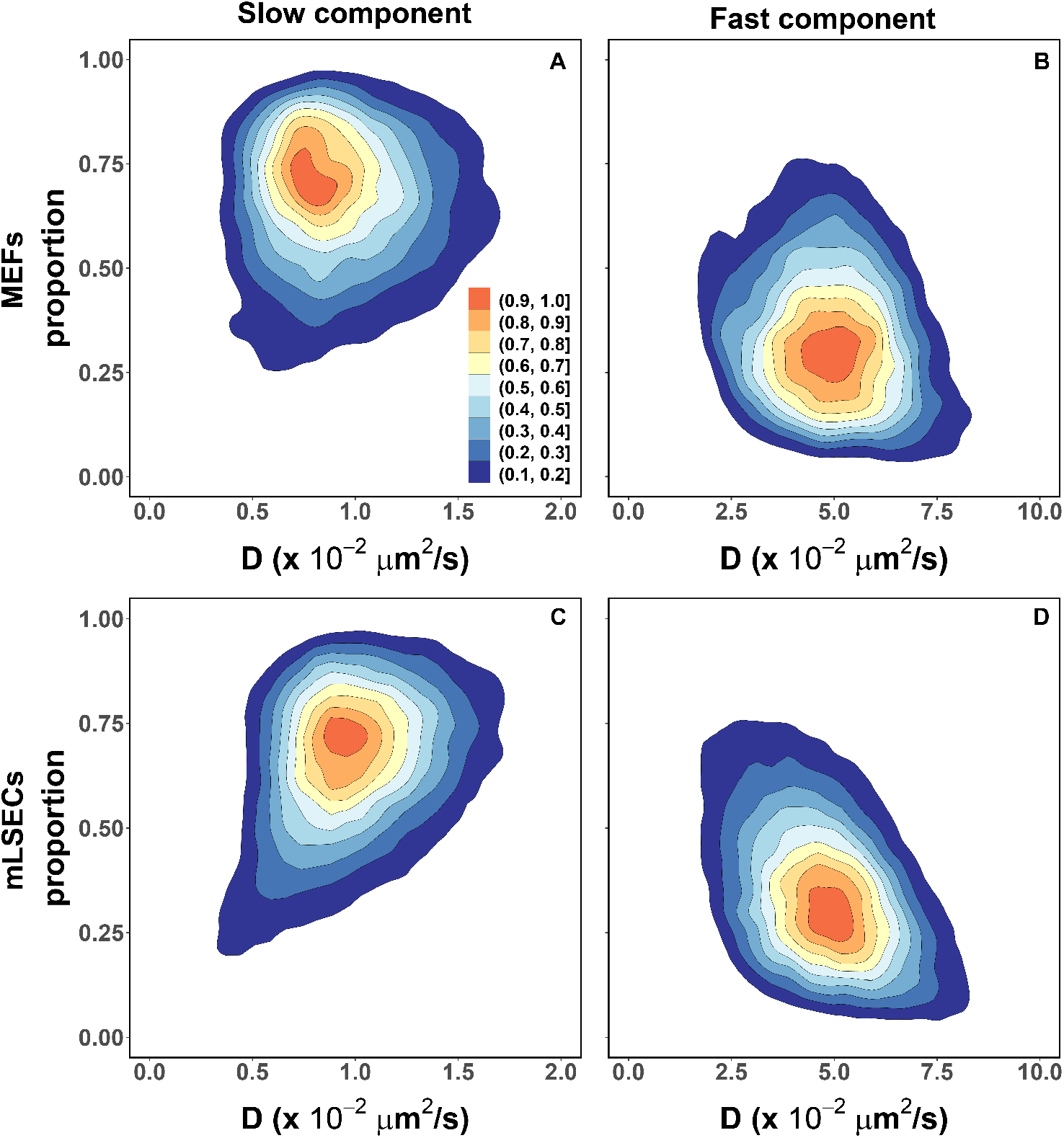
PIEZO1-tdTomato mobile trajectories are heterogeneous. The step-length distribution of most individual mobile trajectories in both (A and B) MEFs (16061 trajectories from 231 MEFs across 23 independent experiments) and (C and D) mLSECs (40611 trajectories from 244 mLSECs across 7 independent experiments) are adequately described by a two-component mixture model (A and C, “slow” component; B and D, “fast” component). The corresponding mixing proportions and apparent diffusion coefficients, D, of each component (shown as joint kernel density estimates) are broadly distributed, and have similar central values, in both cell types. The filled contours are scaled densities.

There is a clear separation between these two mobility states. The “fast” mobility state (Fig. 4B and D) is centered at apparent diffusion coefficient values that are in the same order as previously reported est imates (36, 41). On the other hand, the apparent diffusion coefficients for the “slow” mobility state (Fig. 4A and C) are centered almost one order of magnitude below. Furthermore, for a significant proportion of trajectories (∼62% in MEFs and ∼51% in mLSECs) the “slow” mobility state apparent diffusion coefficient is under the value corresponding to the localization uncertainty (∼1 × 10^−2^ *μ*m^2^/ s, as determined from the trajectory analysis of fixed cells, see Materials and Methods). Despite the diversity in values across trajectories, the parameters of each mobility state are similarly distributed in both cell types, suggesting that the diffusive nature of the PIEZO1-tdTomato mobile class is independent of cell type.

### Manipulation of the lipid membrane composition and modulation of channel activity results in changes to PIEZO1 mobility

Previous studies have shown that changes to membrane composition can affect membrane protein diffusion (37, 58–60). In order to explore the relationship between membrane composition and PIEZO1 mobility, we used chemical agents to manipulate the membrane in MEFs. We then generated and analyzed PIEZO1-tdTomato trajectories from these videos as described above. To assess the effect of the different treatments on the mobility of PIEZO1 puncta, we used the sR_g_-based criterion described above (see Fig. 2) to identify mobile trajectories and computed the odds of observing mobile trajectories upon treatment as the mobile:immobile ratio. The results are shown in Fig. 5 expressed as an odds ratio relative to their corresponding control.

**Figure 5:**
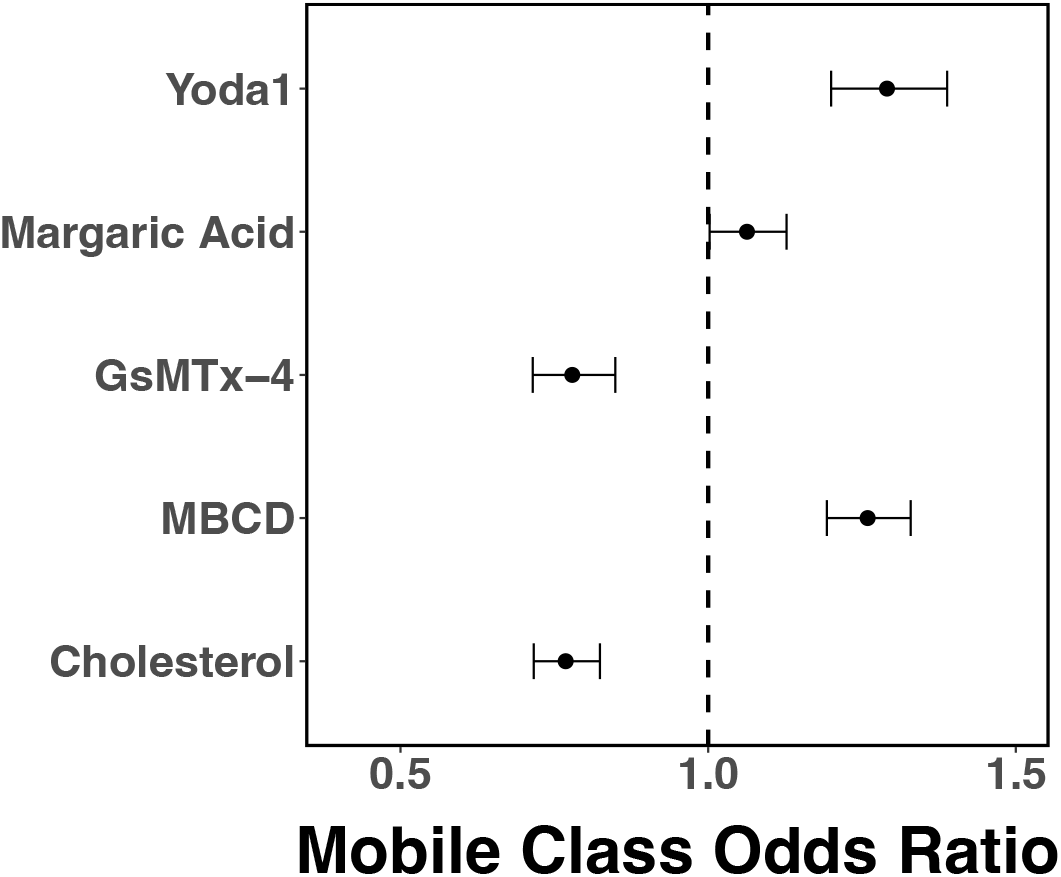
PIEZO1-tdTomato mobile fraction in MEFs is influenced by perturbations to membrane composition and channel activity when compared to the immobile fraction. The odds of observing mobile trajectories increase upon treatment with 10 mM methyl-*β*-cyclodextrin (MBCD, 11097:5666 mobile:immobile trajectories from 86 cells over 3 experiments compared to solvent control, 5906:3670 mobile:immobile trajectories from 66 cells over 3 experiments) or 4 µM Yoda1 (4277:2058 mobile:immobile trajectories from 57 cells over 3 experiments compared to solvent control, 3908:2427 mobile:immobile trajectories sampled from from 185 cells from over 3 experiments), and decrease upon treatment with 100 g/mL cholesterol (3581:2745 mobile:immobile trajectories from 54 cells over experiments compared to 4568:2867 mobile:immobile trajectories from 57 cells over 3 experiments) or 4 µM GsMTx-4 (5048:7506 mobile:immobile trajectories from 99 cells over 3 experiments compared to 1333:1281 mobile:immobile trajectories from 15 cells over 3 experiments). There is no significant effect on the mobile fraction upon treatment with 300 µM margaric acid (5982:3496 mobile:immobile trajectories from 56 cells over 3 experiments compared to solvent control, 5846:3632 mobile:immobile trajectories sampled from from 185 cells from over 3 experiments). The error bars are 95% confidence intervals. Odds ratios upon treatment were computed relative to untreated MEFs (GsMTx-4, MBCD, and cholesterol) or DMSO-treated MEFs (Yoda1 and margaric acid). For contingency tables used in the calculation of the mobile class odds ratio, see Supplementary Tables 1 and 2.

To determine how changes to membrane composition affected PIEZO1 we treated MEFs with 10 mM methyl-*β*-cyclodextrin (MBCD) for 15 minutes in order to deplete cholesterol from the membrane. We chose 10 mM MBCD based on the previous studies in the field ((59, 61)) which examine the impact of cholesterol depletion on membrane protein mobility. Conversely, we next supplemented untreated MEF membranes with 100 µg/mL cholesterol-MBCD for 1 hour in order to simulate the opposite effect on the membrane. In order to verify efficacy of the treatment, we stained the cells with Filipin III, which changes fluorescence upon binding cholesterol (Fig. S1 and S2). We observed lower Filipin III staining in MBCD-treated cells, and higher Filpin III staining in cholesterol-treated cells. Upon MBCD incubation, the odds of observing mobile trajectories increased (Fig. 5). Upon cholesterol supplementation, the odds of observing mobile trajectories decreased. We also incubated cells for 24 hours in 300 µM margaric acid, a fatty acid known to stiffen the membrane (62), in order to explore how membrane stiffness may modify PIEZO1 mobility. Comparison of margaric acid-treated trajectories to the DMSO-treated control trajectories show no significant effects on the mobile proportion (Fig. 5).These results suggest that cholesterol incubation tends to make PIEZO1 less mobile, whereas cholesterol removal increases PIEZO1 mobility. However, margaric acid incubation does not appear to affect PIEZO1 mobility.

We next asked whether the channel’s activation state may affect its mobility. We examined the effect of drugs that modulate PIEZO1 activity on PIEZO1 mobility. GsMTx-4, a spider venom-derived peptide, blocks cation-selective stretch-activated channels and has been shown to inhibit PIEZO1 activity (63). We incubated MEFs in 4 µM GsMTx-4 for 15 min to inhibit PIEZO1 channels. Upon treatment, the odds of observing mobile trajectories are reduced (Fig.5). Thus, GsMTx-4 treatment of PIEZO1 appears to reduce its mobility. We next examined the effect of Yoda1, a chemical activator of PIEZO1, on mobility. We imaged cells treated with 4 µM Yoda1 for 15 minutes, and found an increase in the odds of observing mobile trajectories (Fig.5), suggesting that Yoda1-treated channels are more mobile overall.

We also considered the effect of these treatments on the nature of the mobile class itself. As in the case of untreated MEFs, we found that a two-component mixture model provided an adequate description of the step-length distribution upon treatment for ∼90% of mobile trajectories. Furthermore, although the treatments introduce minor alterations to the most likely values of the mixing proportions and apparent diffusion coefficients, the model parameters’ joint distributions are largely unaltered by the treatments (Fig.6 and Fig. S3).

**Figure 6:**
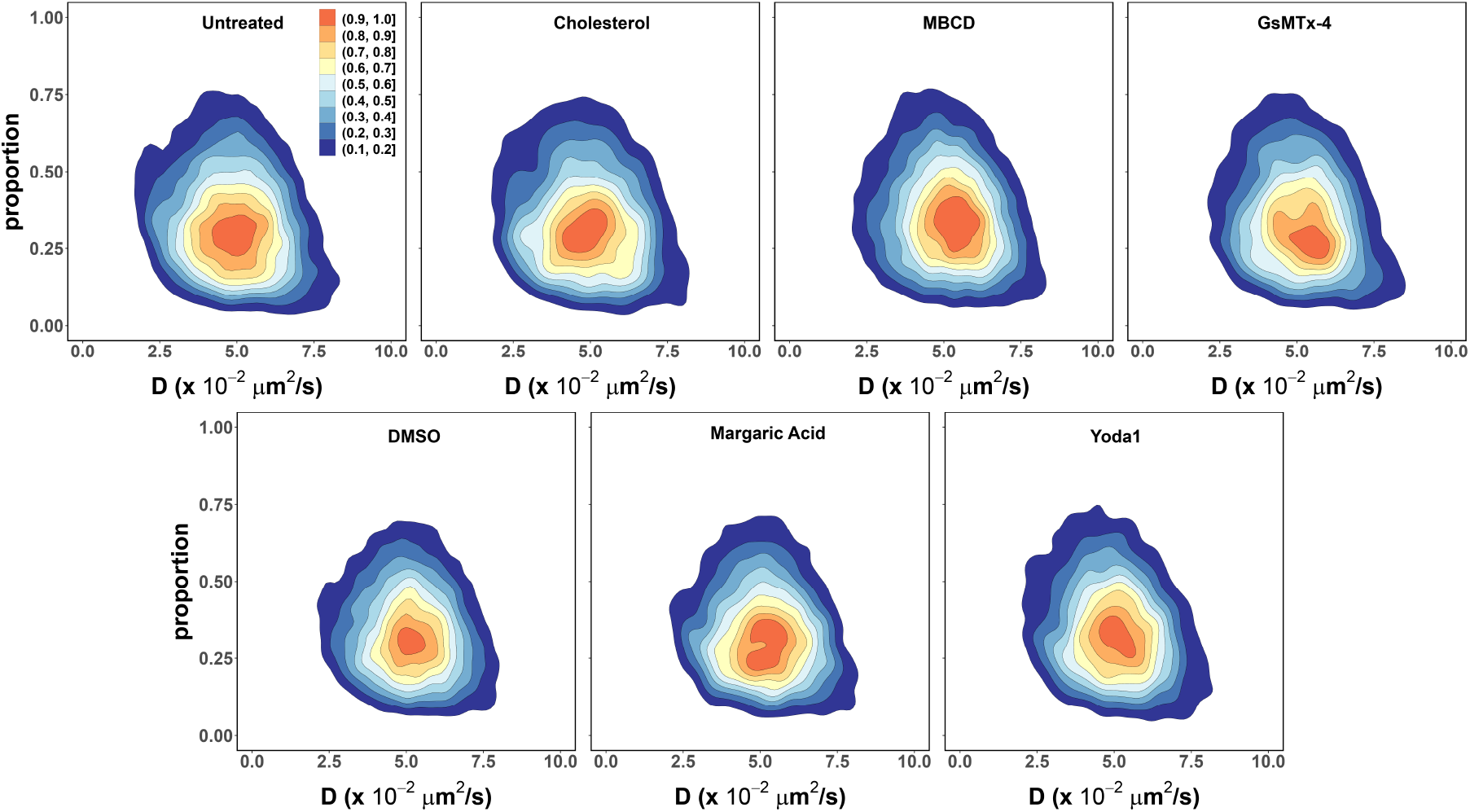
The heterogeneity of PIEZO1-tdTomato mobile trajectories within the “fast” component remains largely unaltered upon treatment. Relative to the corresponding controls (16061 trajectories from 231 untreated MEF cells across 23 independent experiments, 15302 trajectories from 195 DMSO-treated MEF cells across 4 independent experiments), the most likely value of the “fast” component proportion shows minor shifts towards higher values for cholesterol (3565 trajectories from 52 cells across 3 independent experiments), MBCD (15937 trajectories from 128 cells across 10 independent experiments) and Yoda1 (4864 trajectories from 68 cells across 5 independent experiments), and lower values for GsMTx-4 (5035 trajectories from 99 cells across 3 independent experiments), and margaric acid (6109 trajectories from 63 cells across 4 independent experiments), but the overall shape of the parameters’ joint distribution remains the same.

Together, these results indicate that the diffusion of the PIEZO1-tdTomato is sensitive to changes in membrane composition as well as to the activation state of the PIEZO1 channel. On the other hand, the persistent heterogeneity of individual PIEZO1-tdTomato mobile trajectories indicates anomalous diffusion of PIEZO1 in the cell membrane.

### The mobile class is subdiffusive

The available evidence from fluorescence correlation spectroscopy, fluorescence recovery after photobleaching, and SPT indicates that Brownian motion is not the prevalent diffusive behavior of proteins in the membrane environment (reviewed in (39, 64, 65)). Although the mechanistic details of PIEZO1 diffusion in the plasma membrane have yet to be elucidated, our results indicate anomalous subdiffusion of PIEZO1 in membranes.

With the tagged particle’s trajectory denoted as 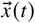, diffusive behavior can be characterized by the so-called time averaged mean squared displacement (TAMSD) (65)

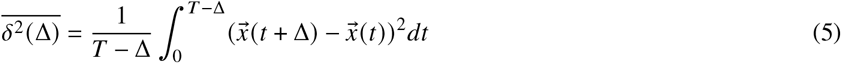

where *T* is the trajectory’s total length in time and Δ is the lag time.

Unrestricted Brownian motion is characterized by a linear time dependence of the TAMSD

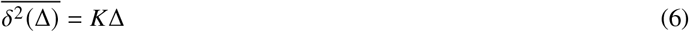

where *K* is a constant. Deviations from this linear behavior, termed anomalous diffusion, are commonly observed in SPT experiments and modeled using a power-law form

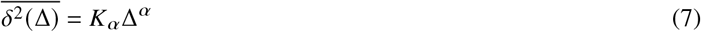

where *α*, the so-called anomalous exponent, is a positive real constant. A time dependence of the TAMSD that is slower than linear (0 < *α* < 1) is called subdiffusion, while a time dependence that is faster than linear (*α* > 1) is called superdiffusion.

We computed the TAMSD of the individual PIEZO1 trajectories in the mobile class (see Fig.7A). Fitting the individual TAMSDs to a power law (Eq. 7) yields a broad distribution of anomalous diffusion exponents, *α* (Fig.7B). We next questioned whether changes to PIEZO1 activity or the membrane composition may impact PIEZO1’s anomalous diffusion. Interestingly, similar results were observed across all the above conditions. These results are not unexpected. SPT is a time limited recording of a stochastic process (Figs. S4 and S5). Therefore, estimates of the anomalous diffusion exponent may vary significantly among trajectories collected from the same experiment. A common practice is to perform an additional average of the TAMSD over an ensemble of *M* collected trajectories,

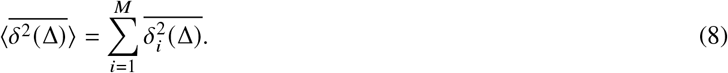

**Figure 7:**
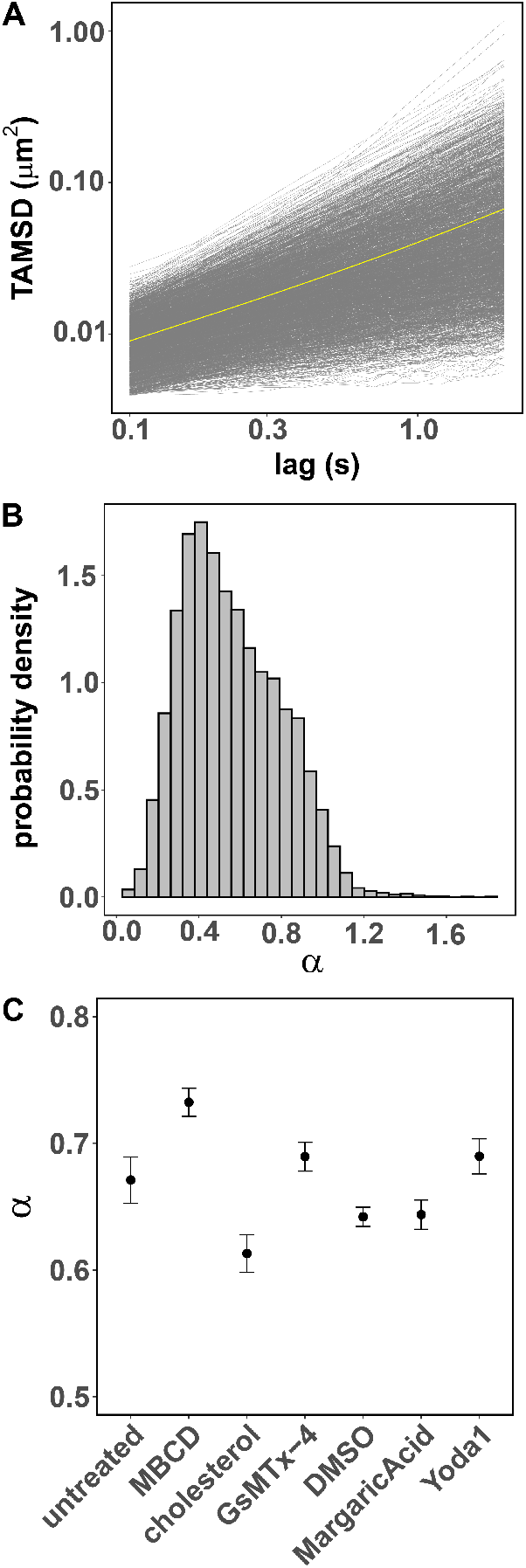
PIEZO1-tdTomato mobile trajectories are subdiffusive. A. The TAMSD as a function of time for single mobile trajectories of PIEZO1-tdTomato expressed in MEFs (a 1% sample of individual trajectories are shown in gray and the ensemble average TAMSD is shown in yellow). B. The power-law exponents (*α*) describing single mobile trajectories TAMSD of PIEZO1-tdTomato expressed in MEFs and mLSECs are broadly distributed. C. The mean estimates of the power-law exponent distributions correct for noise and heterogeneity at the individual trajectory level indicate subdiffusive behavior across all tested conditions. Error bars are 95% confidence intervals (MBCD, cholesterol, and GsMTx-4 treatments using untreated MEFs as control; margaric acid and Yoda1 treatments using DMSO-treated MEFs as control).

The resulting ensemble-averaged TAMSD (EA-TAMSD) (shown in yellow Fig.7A) is sufficient to eliminate the variability associated with time limited measurements and random errors, but it fails to account for measurement noise as well as the intrinsic variability in the particles’ diffusing behavior (49, 66). Accurate estimates of the mean and width of the distribution of anomalous exponents for the ensemble of particles can be obtained from the EA-TAMSD after correcting for these systematic errors as described in (49). The results for the PIEZO1 mobile class indicate a consistent subdiffusive behavior across all the experimental conditions (see Fig.7C).

In contrast to the trajectory spread analysis reported in Fig.5, where changes to the sR_g_ distribution can be directly associated to changes in PIEZO1 mobility upon membrane composition and channel activation state perturbations, an interpretation of the small but statistically significant changes to the anomalous exponent would require a detailed modeling of the diffusion mechanisms, which is beyond the scope of the present TAMSD analysis. Nevertheless, taking together the persistence of anomalous exponent mean estimates well below unity and the results from the two-component mixture model description of the step-length distribution (6) indicate that the heterogeneous nature of individual PIEZO1 mobile trajectories anomalous diffusion is a consistent property of PIEZO1 mobility under a variety of conditions.

## DISCUSSION

Here, we expand upon our previous finding that PIEZO1 channels are mobile (36) by performing single-particle tracking (SPT) of endogenous PIEZO1-tdTomato channel puncta. We observed that PIEZO1 exhibits a heterogeneous diffusive behavior and classified the trajectories into two classes based on their spatial extent - “mobile” and “immobile” using quantitative insights from fixed-cell trajectories. Both classes are present within MEFs, mLSECs, and mNSCs. Analysis of mobile class trajectories from MEFs and mLSECs further demonstrated heterogeneous behavior across trajectories and within trajectories throughout the cell. Taken together, these results indicate non-Brownian diffusion of PIEZO1-tdTomato puncta in the cell membrane.

In order to further probe PIEZO1-tdTomato diffusion, we manipulated cellular membrane composition and channel activity using chemical reagents. We consistently observed “mobile” and “immobile” trajectories even when cells were treated with drugs that manipulate membrane composition (cholesterol, MBCD, and margaric acid) and PIEZO1 activity (Yoda1 and GsMTx-4). Furthermore, channel activation via Yoda1 and cholesterol removal via MBCD increased the odds of observing PIEZO1-tdTomato mobile trajectories relative to the respective controls. Conversely, we found that channel inhibition via GsMTx-4 and supplementation of the membrane with cholesterol decreased the odds of observing mobile trajectories. Treatment with margaric acid, which stiffens the membrane and inhibits PIEZO1 (62), has no statistically significant effect on the odds of observing mobile trajectory. In every case, however, mobile trajectories remain heterogeneous and subdiffusive.

Interestingly, we observed PIEZO1-tdTomato puncta with low or no mobility that aggregated in structures reminiscent of focal adhesions (Fig. 1A, green inset). In cells with contractile myosin IIA, PIEZO1 has previously been found to localize to focal adhesions in human foreskin fibroblasts, resulting in integrin *β*3 adhesion disassembly and turnover (53). In cells without myosin IIA expression, PIEZO1 was dispersed throughout the cell. Another study demonstrated that PIEZO1 localizes at focal adhesions to trigger integrin-FAK signaling and tissue stiffening in human gliomas (54). We previously found that PIEZO1 in adherent cells is activated by cell-generated traction forces, which are transmitted to the substrate at focal adhesions (19, 36). PIEZO1 is more active at cellular regions of high traction forces than at regions of low traction forces. In this scenario, PIEZO1 may localize to focal adhesions to better sense and transduce these cell-generated mechanical forces. Together, these studies suggest that components in focal adhesions may have a role in modulating PIEZO1 mobility and activity.

In this study, we observed multiple mobile PIEZO1 populations throughout the cell in different cell types. Our reports complement findings from previous PIEZO1 studies that examine PIEZO1 mobility. Ridone *et al*. heterologously expressed PIEZO1 tagged with GFP (PIEZO1-GFP). Using TIRF microscopy and the ensemble-level technique, spatiotemporal k-space image correlation spectroscopy, they characterized the mobility of PIEZO1-GFP (37). When they depleted cholesterol from the membrane using MBCD, they observed that PIEZO1 clusters in the membrane were disrupted, and that PIEZO1-GFP diffusion rates were increased. Similar to their findings, we observe that PIEZO1-tdTomato mobility is increased upon MBCD treatment. Vaisey *et al*. examined endogenously expressed PIEZO1 tagged with a hemagglutinin (HA) tag (PIEZO1-HA) in red blood cells. They reported mobile PIEZO1 trajectories distributed on the red blood cell surface and particularly concentrated in the red blood cell “dimple” (41). Of note, they observed a heterogeneity in PIEZO1 diffusors. Consistent with their observations, we also found heterogeneous behavior of mobile PIEZO1.

By examining PIEZO1 trajectories generated through the SPT of individual puncta, we identified two populations of PIEZO1 - a “mobile” class and an “immobile” class. Other membrane proteins have also been shown to display mobile and immobile fractions in the plasma membrane. ORAI1, an ion channel that allows extracellular Ca^2+^ influx upon internal Ca^2+^ store depletion, was fluorescently labeled with mCherry (ORAI1-mCherry), and its mobility was classified into four classes: directed, linear, confined, and transiently confined (59). Similarly, the mobility of *α*PS2C*β*PS integrin, tracked with quantum dots and SPT, was classified into three classes (non-confined diffusion, confined diffusion, and immobility) (67). Glycine receptors in spinal cord neurons, tracked with BODIPY-strychnine, also demonstrated mobile and confined fractions when observed via Fluorescence Recovery After Photobleaching (68). Cystic fibrosis transmembrane conductance regulator (CFTR), tagged with GFP (CFTR-GFP), was classified into two classes: confined and unconfined (60). MEC-2, a stomatin-like protein that is a part of channel complex responsible for touch sensation in *C. elegans*, has also been observed in distinct populations within touch receptor neurons: static, immobile puncta along the neurite, highly mobile puncta in the cell body, and puncta displaying directed motion from the cell body to the distal neurite (69). The immobile class that we observe in PIEZO1-tdTomato mobility could stem from transient interactions with components of the membrane or the cytoskeleton. One possibility is that the actin cytoskeleton could be acting as barriers to PIEZO1 diffusion (70). Alternatively, PIEZO1-tdTomato could be immobilized within lipid microdomains, thereby limiting the mobility of these proteins. Taken together, these studies indicate that transmembrane proteins display elaborate mobility behaviors, which likely reflects the complexity of the plasma membrane and the associated cytoskeleton.

In our study, we manipulate the membrane composition to observe how these changes affect PIEZO1-tdTomato mobility. We find that cholesterol-depleted membranes have a higher likelihood of “mobile” PIEZO1-tdTomato, whereas cholesterol-supplemented membranes have a lower likelihood of “mobile” PIEZO1-tdTomato. Other groups have examined the effect of changing membrane composition on the diffusion of other transmembrane proteins, and reported similar results to our findings. Notably, Ridone *et al*. observed an increase in PIEZO1’s diffusion constant when cholesterol was depleted using MBCD, further supporting our findings. Curiously, when Ridone *et al*. incubated their cells in cholesterol, they did not observe a significant shift in mobility. We similarly do not see an appreciable change in the mobile fraction upon cholesterol supplementation (Fig.6). However, when we calculated the mobile class odds ratio, we observed a decrease in the likelihood of mobile PIEZO1 (Fig. 6), suggesting that cholesterol supplementation affects PIEZO1 mobility by changing the partitioning between mobile and immobile classes. CFTR-GFP-expressing cells treated with cholesterol oxidase to deplete cholesterol demonstrated a decreased confined fraction (60), an effect that could be reversed upon cholesterol supplementation. ORAI1-mCherry was found to be more mobile following MBCD treatment, also consistent with our results (59). Serotonin transporters tagged with quantum dots (71) and dopamine transporters tagged with yellow fluorescent protein (61) also demonstrated an increase in diffusion following MBCD treatment. These studies support our observations: cholesterol depletion increases the mobility of confined transmembrane proteins, and cholesterol supplementation decreases the mobility of these molecules. Interestingly, PIEZO1-tdTomato in MEFs incubated in margaric acid appeared to have no significant shifts in mobility, a counterintuitive result given margaric acid’s role in stiffening the membrane and inhibiting the channel (62). This unexpected result suggests that the effects of margaric acid on PIEZO1 may be complex, and that this intriguing result warrants further study.

At the individual trajectory level, we observed that the heterogeneity within the mobile PIEZO1-tdTomato class is evident in two different cell types, and it persists in MEFs exposed to different drug treatments, suggesting that this may be a fundamental characteristic of PIEZO1 mobility in the plasma membrane. Moreover, we observe that PIEZO1-tdTomato is subdiffusive across all conditions studied; however, further study is required to understand the nature and origin of PIEZO1 subdiffusion and heterogeneity, including the extent to which they rely on the membrane environment, channel clustering, and channel gating states. One potential explanation for these results is that it could originate from changes in oligomerization. For example, stomatin-like protein-3 (STOML3). STOML3, a cholesterol-binding protein that oligomerizes, has been shown to tune PIEZO1 mechanosensitivity (72, 73). Thus, it is possible that STOML3 also clusters PIEZO1 into cholesterol-rich domains, and MBCD treatment disrupts these PIEZO1-STOML3-cholesterol clusters.

PIEZO1 mobility may have several important implications. At the physiological level, we recently showed the importance of dynamic relocalization of the channel in cell migration and wound healing. In non-migrating cells, PIEZO1 is distributed randomly on the cell surface. In single, migrating keratinocytes (18), we found an accumulation of PIEZO1, organized in macroclusters at the cell rear, which modulates rear retraction and thereby the speed of cell migration. In keratinocyte monolayers, we observed similar PIEZO1 enrichment at regions of the wound edge (18) that resulted in local retraction, reducing the rate of monolayer migration and wound healing. Thus, dynamic shifts in PIEZO1 localization and clustering are physiologically important, highlighting the need to pursue further SPT studies of PIEZO1, and expanding them to encompass longer timeframes in cells transitioning from stationary to migrating.

At the subcellular level in non-migrating cells, it is possible that PIEZO1 mobility may enable fewer channels to explore a larger domain of the cell, allowing the channel to more efficiently transduce mechanical forces. PIEZO1 mobility may also function as a mechanism to dynamically adjust cellular response to mechanical forces. Mechanical forces can act upon a cell at any time, from anywhere, and PIEZO1 mobility may allow the channel to move towards or away from mechanical stimuli. Open and closed channels may exhibit different mobilities, allowing the cell to modulate mechanotransduction. For instance, closed channels may be more mobile than open channels, as observed in TRPV1 activated with capsaicin (74). This would allow closed-mobile channels to explore the cell in search of mechanical cues, and for open channels to linger at cellular regions experiencing mechanical stimuli. Conversely, open channels may be more mobile than closed channels. In this case, open-mobile channels may explore the cell, and may move towards or away from mechanical stimuli. If these channels localize toward mechanical stimuli, they can better engage with mechanical forces. Alternatively, open channels may venture away from mechanical stimuli, thereby terminating mechanotransduction. Channel mobility and its relationship to mechanical stimuli likely involves complex interactions between the channel, membrane, and cytoskeleton that remains to be explored. Our findings set the stage for future work examining PIEZO1 mobility in the context of channel activity and its physiological roles.

## AUTHOR CONTRIBUTIONS

M.M.P., D.J.T., and J.A.F. conceptualized the research. A.T.L., J.A.F., G.D.D., G.A.B, E.L.E. were involved with the methodology. A.T.L., J.A.F. were responsible for data curation, investigation, formal analysis, and validation. M.M.P. and D.J.T. were responsible for overseeing the project, and for funding acquisition and resources. A.T.L., J.A.F, D.J.T., and M.M.P wrote the manuscript. A.T.L. and J.A.F. contributed equally to this work. All authors reviewed and edited the manuscript.

## ACKNOWLEDGMENTS

We thank Dr. Ardem Patapoutian for the gift of the PIEZO1-tdTomato mice, Dr. Vivek Tyagi for contributions during the early stages of the manuscript, and Michael Thanh-Phong Vu for assistance with trajectory generation from TIRFM videos, and the Pathak and Tobias groups for invaluable discussions. This work utilized computing resources operated by the Research Cyberinfrastructure Center at the University of California, Irvine.

## SUPPLEMENTAL MATERIAL

The code used for the trajectory analysis is available on GitHub at https://github.com/Pathak-Lab/Piezo1-tdTomato-Trajectory-Ana

**Supplemental Figure 1:**
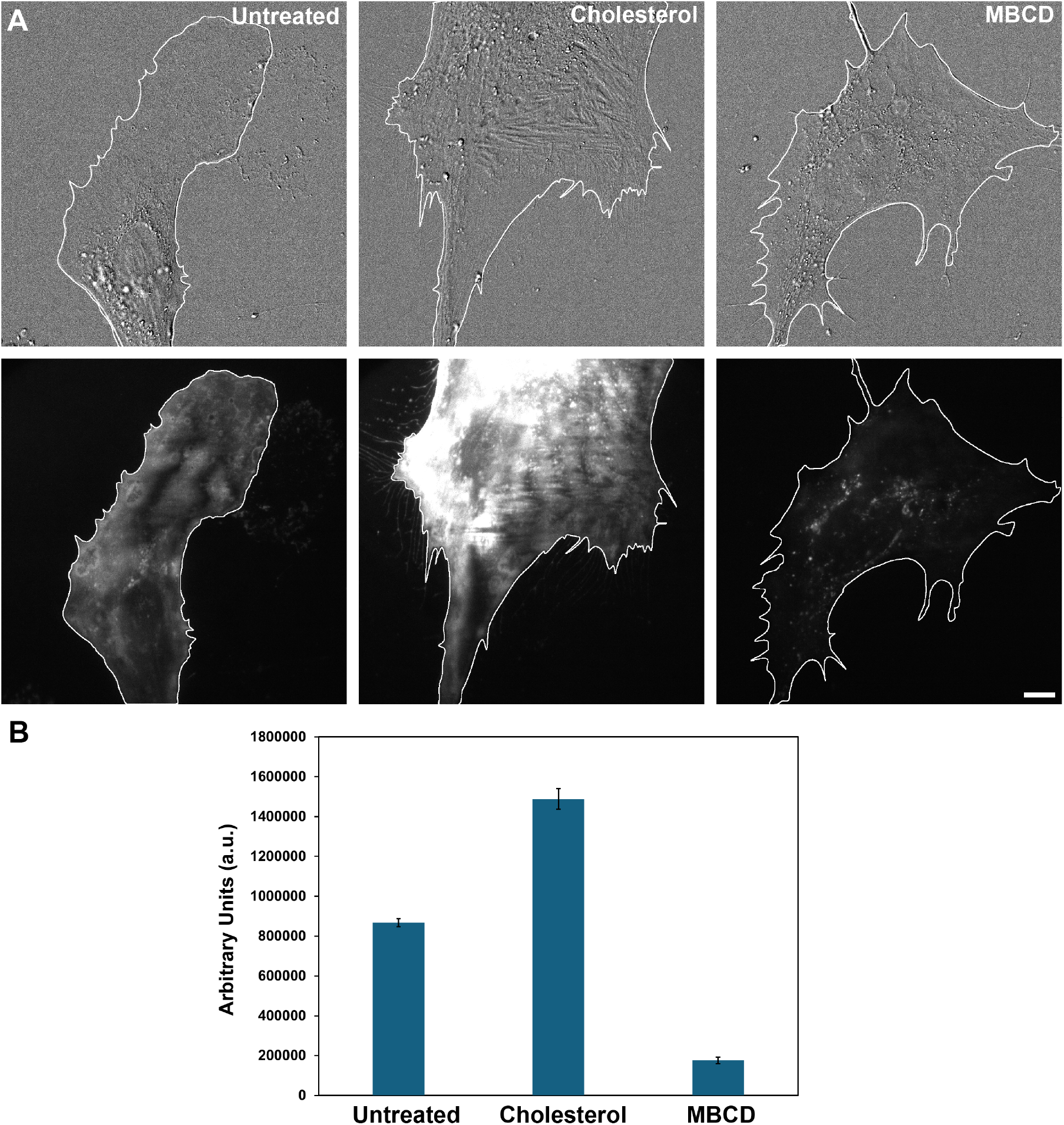
Cholesterol supplementation via cholesterol incubation and cholesterol reduction via MBCD increases and decreases cholesterol signal respectively as determined using Filipin III. A. Representative DIC and TIRF microscopy images of PIEZO1-tdTomato MEFs stained for free cholesterol using Filipin III in the Untreated, Cholesterol-treated, and MBCD-treated conditions respectively. Cell boundary is denoted by white line. TIRFM images were brightness contrast adjusted the Untreated condition. Scale bar = 10 µm. B. Mean calculated total cell fluorescence (CTCF) of Filipin III fluorescence. Intensity calculations were performed using Fiji (ImageJ). Untreated control (n=15), Cholesterol-treated (n=16), MBCD-treated (n=15). Units are in arbitrary units (a.u.). Error bars represent mean CTCF ± SEM.

**Supplemental Figure 2:**
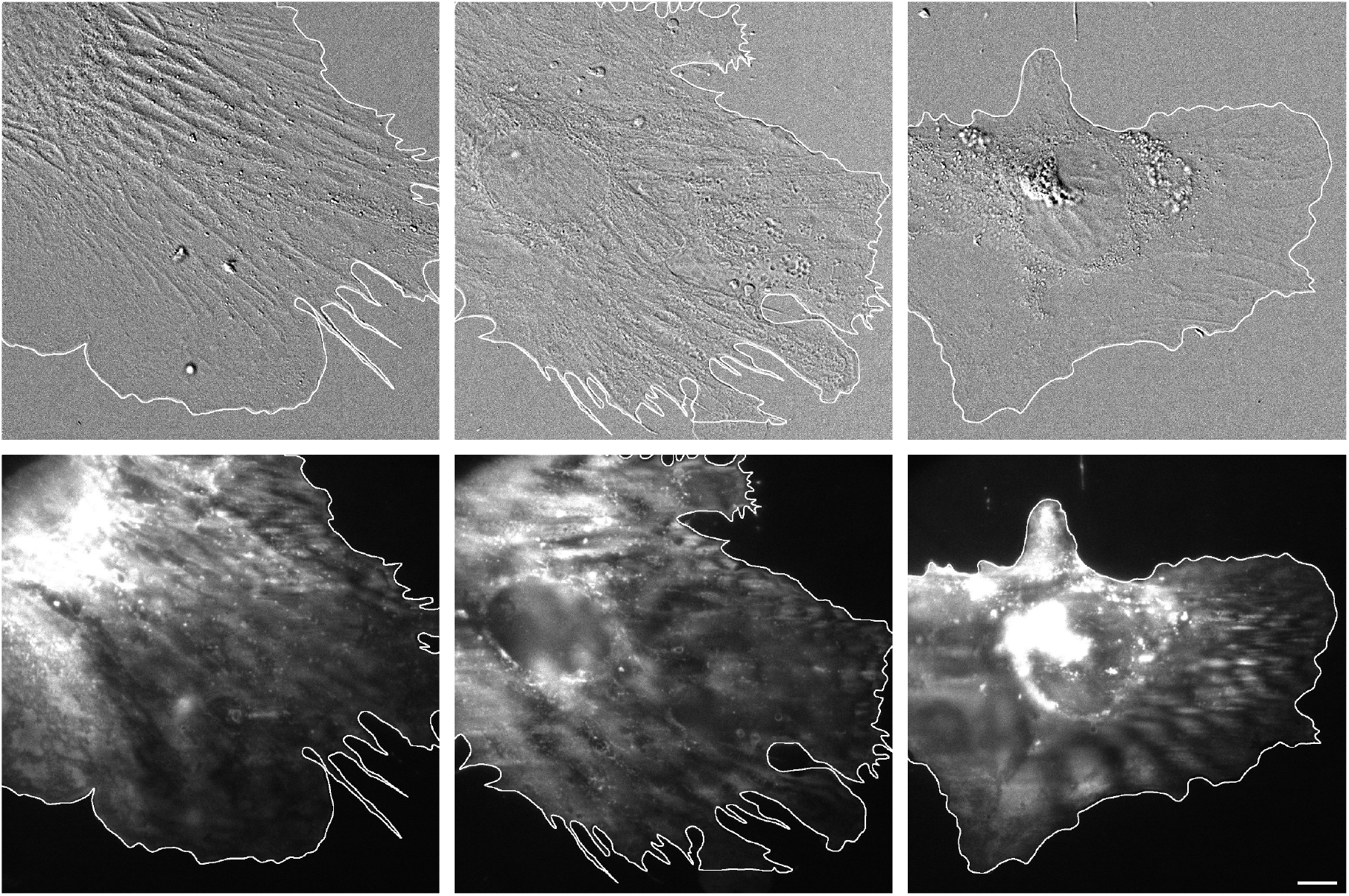
Representative DIC and TIRF microscopy images of PIEZO1-tdTomato MEFs stained for free cholesterol with Filipin III in the Cholesterol-treated condition. Cell boundary is denoted by white line. TIRFM images were brightness contrast adjusted the Untreated condition. Scale bar = 10 µm.

**Supplemental Table 1:** Contingency table used to calculate the mobile class odds ratio for drugs that can be compared with untreated control MEFs. All conditions were obtained over 3 experiments.

|  | Mobile | Immobile |
| --- | --- | --- |
| MBCD | 11097 | 5666 |
| MBCD control | 5906 | 3670 |
| Cholesterol | 3581 | 2745 |
| Cholesterol control | 4568 | 2867 |
| GsMTx-4 | 5048 | 7506 |
| GsMTx-4 control | 1333 | 1281 |

**Supplemental Table 2:**
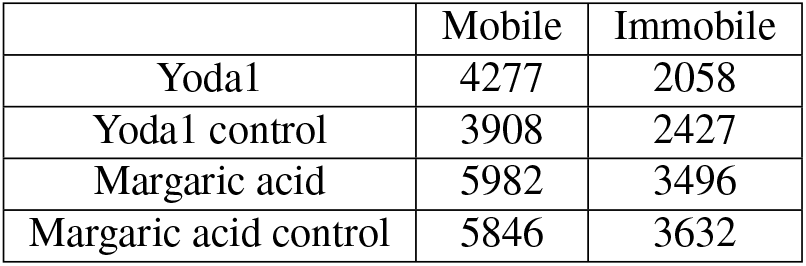
Contingency table used to calculate the mobile class odds ratio for drugs that can be compared with DMSO-treated MEFs. All conditions were obtained over 3 experiments.

**Supplemental Figure 3:**
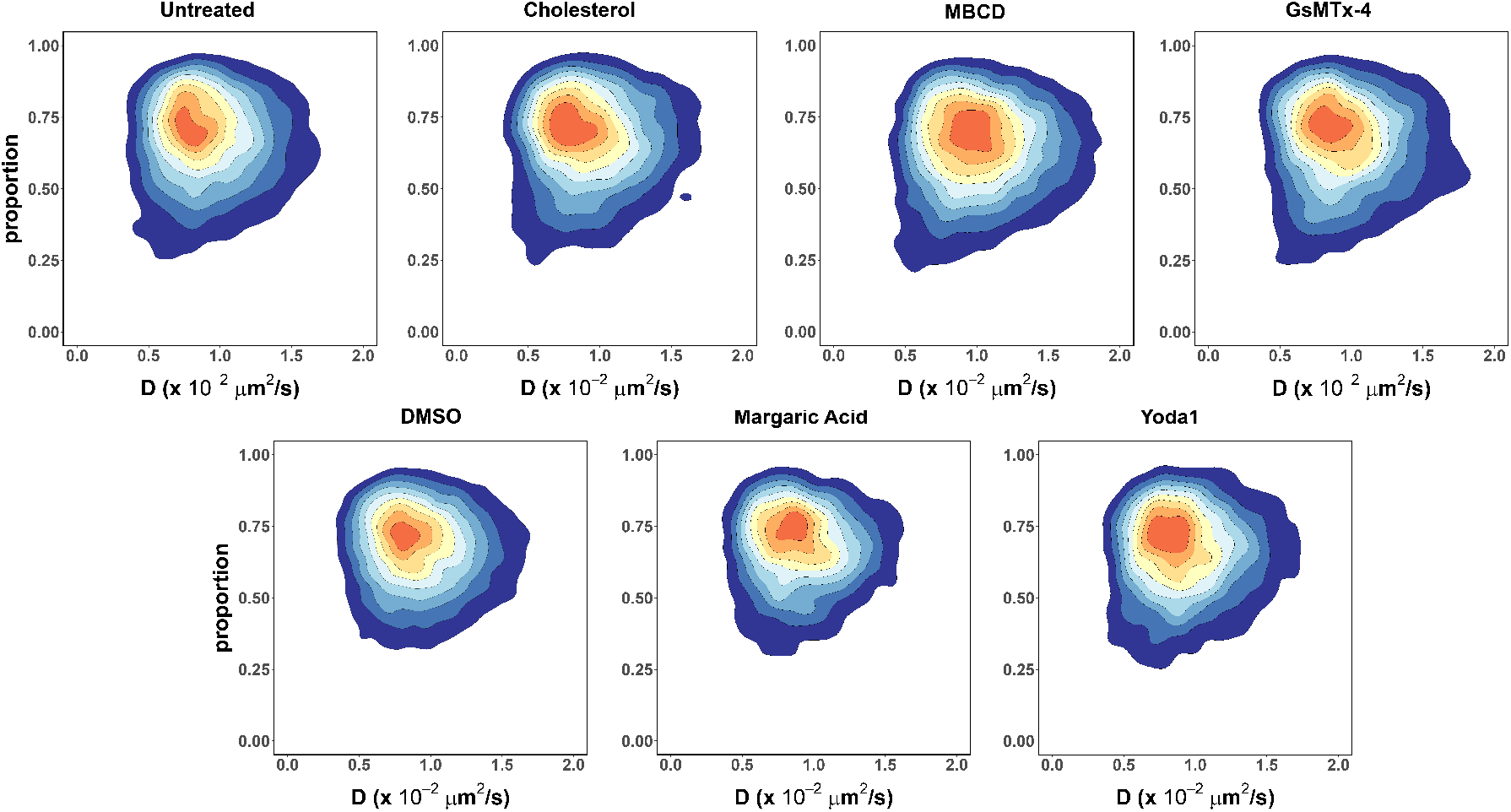
The heterogeneity of PIEZO1-tdTomato immobile trajectories of the “slow” component remains largely unaltered upon treatment. The overall shape of the joint distribution of the parameters remains unchanged compared to the corresponding controls.

**Supplemental Figure 4:**
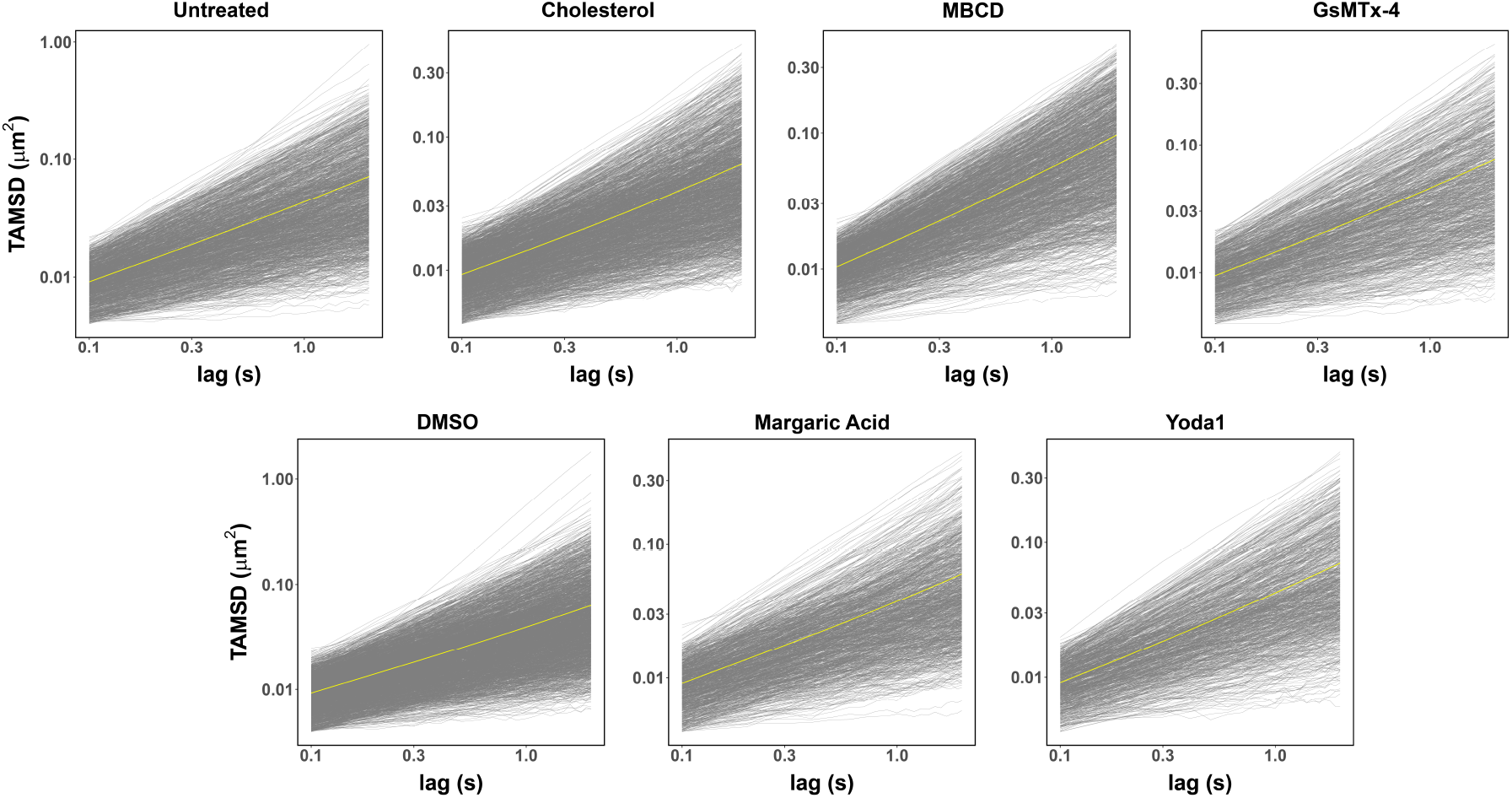
The TAMSD as a function of time for single mobile trajectories of PIEZO1-tdTomato expressed in MEFs and treated with assorted drugs (a 1% sample of individual trajectories are shown in gray and the ensemble average TAMSD is shown in yellow).

**Supplemental Figure 5:**
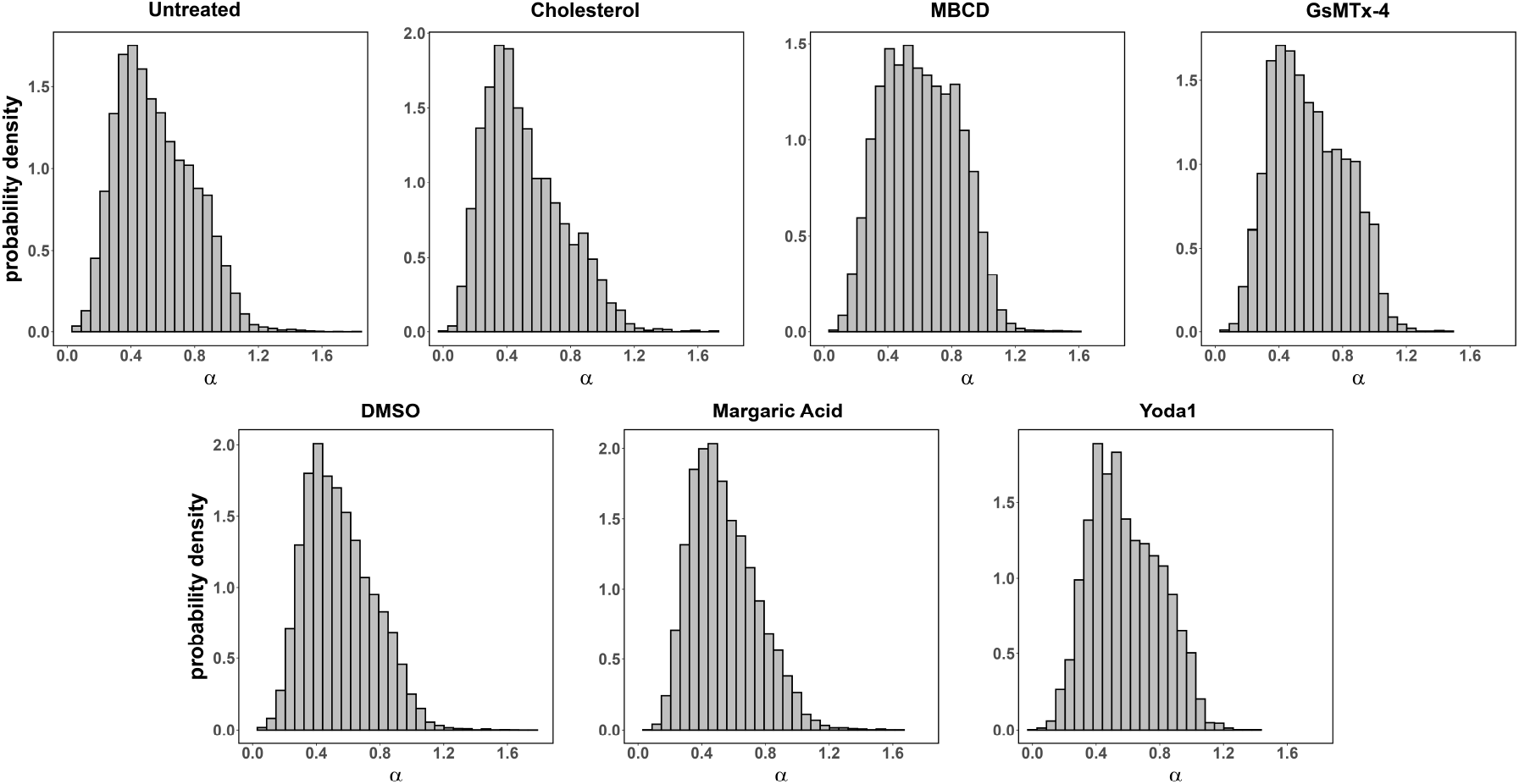
The power-law exponents (*α*) describing single mobile trajectories TAMSD of PIEZO1-tdTomato expressed in MEFs treated with cholesterol, MBCD, GsMTx-4, margaric acid, Yoda1, and their respective controls are broadly distributed.

## SUPPLEMENTARY VIDEOS

Supplementary Video 1: TIRF microscopy reveals heterogeneity in PIEZO1-tdTomato mobility. A. Representative TIRF image of PIEZO1-tdTomato puncta in live MEFs harvested from PIEZO1-tdTomato reporter mice. The white line denotes the cell boundary. Insets show enlarged regions of interest. The green inset is representative of regions where puncta appear mobile, whereas the blue inset is representative of regions where puncta show little or no mobility.

Scale bars = 10 µm.

Supplementary Video 2: TIRF microscopy videos of PIEZO1-tdTomato puncta from zoomed-in regions of live (left) and paraformaldehyde-fixed (right) MEFs. Trajectories are overlaid on to the video. Note the greater mobility of puncta in the live cell compared to fixed cell.

Scale bar = 1 µm.

## SUPPLEMENTAL METHODS: VALIDATION OF TRAJECTORY GENERATION PARAMETERS

In order to guide and validate our trajectory generation parameters, we conducted several additional analyses, which we have detailed in this Supplemental Methods section. Prior to any analysis, we needed to select the ideal pixel linking distance between puncta. When linking trajectories, ID switching, a process in which puncta from different trajectories can be incorrectly linked, can occur when the pixel linking distance between puncta is too large. In this case, multiple trajectories could be combined into a larger trajectory, resulting in inaccurate trajectory measurements. On the other hand, trajectory fragmentation can occur when the pixel linking distance is too small. In this situation, trajectories generated will be incomplete and truncated. Thus, generating trajectories in Single Particle Tracking (SPT) studies involves an inherent trade-off between these two situations. These technical tests were performed on a small subset of our data (three representative experimental recordings, which we will now refer to as “test data” in this section) in order to guide the parameters used to generate trajectories.

Based on visual inspection of the data, we first estimated that a three-pixel linking distance would be reasonable for trajectory generation. To test this choice, we measured the nearest neighbor (NN) distance between each linked particle and other particles in the image frame. The KDTree function from the scikit-learn library (75) was used to efficiently search for the NN to each punctum in a trajectory and the Euclidean distance was recorded (in pixels). This process was repeated for all frames in a single recording and the data summarized by the respective mean of each trajectory class (mobile or immobile). The resulting mean NN-distance values averaged over three separate recordings are shown in Supp. Methods Fig. 1A. In both trajectory classes, the mean NN-distance value is larger than the three-pixel distance used for linkage.

To quantify the displacement of linked puncta between consecutive frames, we computed the mean step length for each trajectory, given for an *N*-point trajectory, 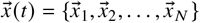, by ^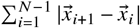^. The calculations were performed using Flika (43) and a custom Python script. Similar to the NN distance calculation, we summarized the data from each recording by taking the means over each trajectory class. The resulting average values obtained from our test data are shown in shown in Supp. Methods Fig. 1B. In both trajectory classes, the mean step length is shorter than the three-pixel distance used for linkage, suggesting that we were able to successfully link most of the trajectories properly with a three-pixel linking distance.

**Supplemental Methods Figure 1:**
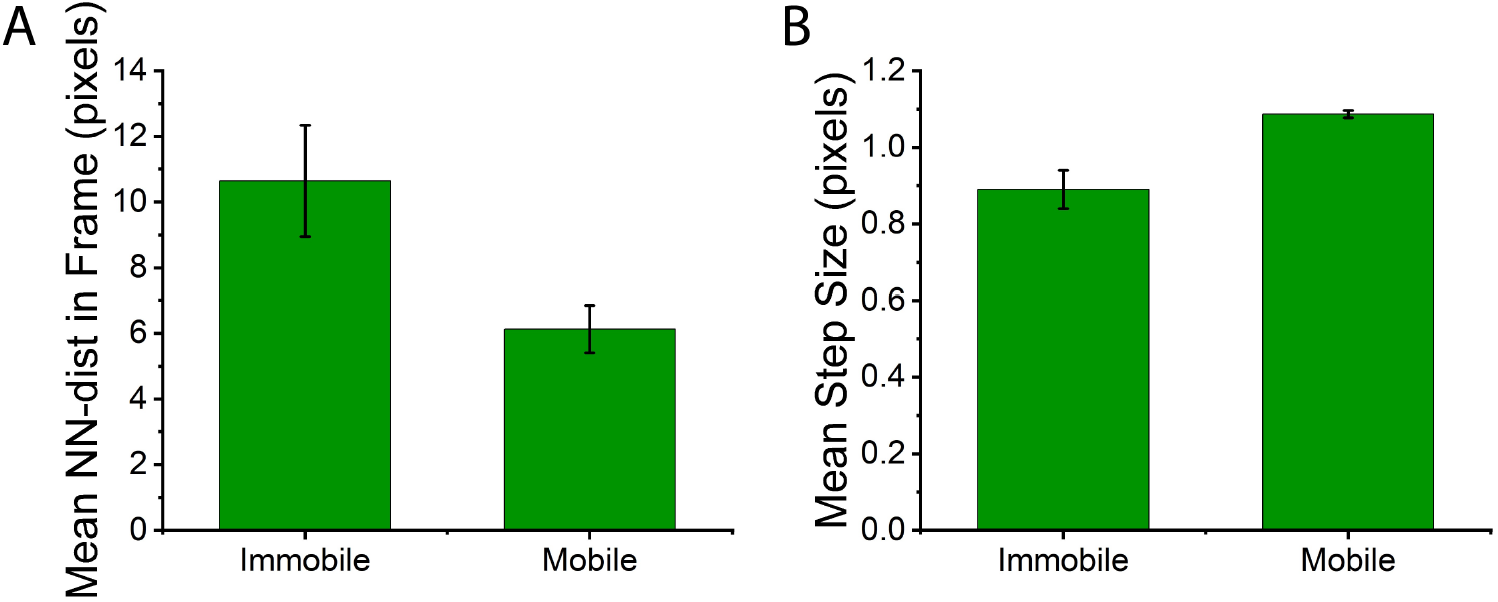
(A) Mean distance to the nearest-neighbor (NN) particle in a recording frame for each punctum in a trajectory. The mean NN distance between puncta in a recording frame for both immobile and mobile trajectories is larger than the three-pixel linking distance. (B) Mean step length between linked puncta in a trajectory. Trajectories from both (A) and (B) were generated using Flika and classified as mobile or immobile based on their s R_g_ values as described in Materials and Methods. The average of mean values from three 20 s recordings are shown, and the error bars are standard deviations.

We considered next the effect of the linkage distance values on trajectory classification. Using two second recordings from our test data, we implemented a range of pixel linking (1-20 pixels), which were generated using Flika (43) and a custom python script for iterating through the different parameters (Supp. Methods Fig. 2). Classification was performed as described in the manuscript. The two second recordings were truncated from the full-length recordings used in the manuscript to reduce the amount of computation required, which otherwise would have been restrictive.

We observed an increase in the proportion of mobile trajectories from 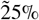 at a one-pixel linking distance to 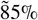 at fifteen pixels and above, suggesting false-linkages between trajectories at higher pixel linkages and resulting in a classification of every trajectory as mobile. We similarly noted that the opposite effect occurred for immobile trajectories, hinting that these trajectories were falsely truncated at a one-pixel threshold. Thus, the three-pixel threshold (dashed blue line in Supp. Methods Fig. 2) provided a good balance for capturing correct vs false links.

**Supplemental Methods Figure 2:**
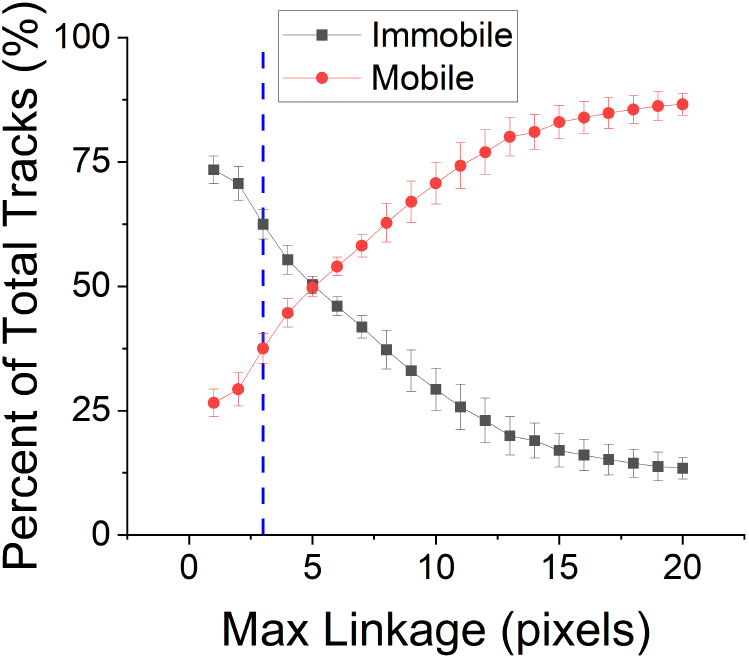
The effect of linkage distance value on trajectory classification. Trajectories were established for 2 s test data using a range of linkage parameters (1-20 pixels). The plot shows mean values over the three recordings. The error bars are standard deviations. The dashed blue line is indicative of 3-pixel linkage, which we use in our analyses.

While a three-pixel cutoff may result in some truncated trajectories and an underestimation of actual trajectory mobility, this does not affect the main findings of the paper as the majority of step sizes fall below three pixels in the mobile class. Together, these analyses demonstrate that the three-pixel threshold provides a good balance between ID switching and fragmented trajectories.

## REFERENCES

1. Martinac, B., T. Nomura, G. Chi, E. Petrov, P. R. Rohde, A. R. Battle, A. Foo, M. Constantine, R. Rothnagel, S. Carne, E. Deplazes, B. Cornell, C. G. Cranfield, B. Hankamer, and M. J. Landsberg, 2014. Bacterial Mechanosensitive Channels: Models for Studying Mechanosensory Transduction. Antioxidants &amp; Redox Signaling 20:952–969. 10.1089/ars.2013.5471.

2. Murthy, S. E., A. E. Dubin, T. Whitwam, S. Jojoa-Cruz, S. M. Cahalan, S. A. R. Mousavi, A. B. Ward, and A. Patapoutian, 2018. OSCA/TMEM63 are an evolutionarily conserved family of mechanically activated ion channels. eLife 7. 10.7554/elife.41844.

3. Poole, K., 2022. The Diverse Physiological Functions of Mechanically Activated Ion Channels in Mammals. Annual Review of Physiology 84:307–329. 10.1146/annurev-physiol-060721-100935.

4. Sukharev, S. I., P. Blount, B. Martinac, F. R. Blattner, and C. Kung, 1994. A large-conductance mechanosensitive channel in E. coli encoded by mscL alone. Nature 368:265–268. 10.1038/368265a0.

5. Driscoll, M., and M. Chalfie, 1991. The mec-4 gene is a member of a family of Caenorhabditis elegans genes that can mutate to induce neuronal degeneration. Nature 349:588–593. 10.1038/349588a0.

6. Coste, B., J. Mathur, M. Schmidt, T. J. Earley, S. Ranade, M. J. Petrus, A. E. Dubin, and A. Patapoutian, 2010. Piezo1 and Piezo2 Are Essential Components of Distinct Mechanically Activated Cation Channels. Science 330:55–60.

7. Ranade, S. S., Z. Qiu, S.-H. Woo, S. S. Hur, S. E. Murthy, S. M. Cahalan, J. Xu, J. Mathur, M. Bandell, B. Coste, Y.-S. J. Li, S. Chien, and A. Patapoutian, 2014. Piezo1, a mechanically activated ion channel, is required for vascular development in mice. Proc. Natl. Acad. Sci. U.S.A. 111:10347–10352.

8. Li, J., B. Hou, S. Tumova, K. Muraki, A. Bruns, M. J. Ludlow, A. Sedo, A. J. Hyman, L. McKeown, R. S. Young, N. Y. Yoldasheva, Y. Majeed, L. A. Wilson, B. Rode, M. A. Bailey, H. R. Kim, Z. Fu, D. A. L. Carter, J. Bilton, H. Imrie, P. Ajuh, T. N. Dear, R. M. Cubbon, M. T. Kearney, K. R. Prasad, P. C. Evans, J. F. X. Ainscough, and D. J. Beech, 2014. Piezo1 integration of vascular architecture with physiological force. Nature 515:279–282.

9. Nonomura, K., V. Lukacs, D. T. Sweet, L. M. Goddard, A. Kanie, T. Whitwam, S. S. Ranade, T. Fujimori, M. L. Kahn, and Patapoutian, 2018. Mechanically activated ion channel PIEZO1 is required for lymphatic valve formation. Proc. Natl. Acad. Sci. U.S.A. 115:12817–12822.

10. Sun, W., S. Chi, Y. Li, S. Ling, Y. Tan, Y. Xu, F. Jiang, J. Li, C. Liu, G. Zhong, D. Cao, X. Jin, D. Zhao, X. Gao, Z. Liu, B. Xiao, and Y. Li, 2019. The mechanosensitive Piezo1 channel is required for bone formation. eLife 8:e47454.

11. Zeng, W.-Z., K. L. Marshall, S. Min, I. Daou, M. W. Chapleau, F. M. Abboud, S. D. Liberles, and A. Patapoutian, 2018. PIEZOs mediate neuronal sensing of blood pressure and the baroreceptor reflex. Science 362:464–467.

12. Hill, R. Z., M. C. Loud, A. E. Dubin, B. Peet, and A. Patapoutian, 2022. PIEZO1 transduces mechanical itch in mice. Nature 607:104–110.

13. Ranade, S. S., S.-H. Woo, A. E. Dubin, R. A. Moshourab, C. Wetzel, M. Petrus, J. Mathur, V. Bégay, B. Coste, J. Mainquist, J. Wilson, A. G. Francisco, K. Reddy, Z. Qiu, J. N. Wood, G. R. Lewin, and A. Patapoutian, 2014. Piezo2 is the major transducer of mechanical forces for touch sensation in mice. Nature 516:121–125.

14. Chesler, A. T., M. Szczot, D. Bharucha-Goebel, M. Čeko, S. Donkervoort, C. Laubacher, L. H. Hayes, K. Alter, C. Zampieri, C. Stanley, A. M. Innes, J. K. Mah, C. M. Grosmann, N. Bradley, D. Nguyen, A. R. Foley, C. E. Le Pichon, and C. G. Bönnemann, 2016. The Role of PIEZO2 in Human Mechanosensation. N. Engl. J. Med. 375:1355–1364.

15. Woo, S.-H., V. Lukacs, J. C. de Nooij, D. Zaytseva, C. R. Criddle, A. Francisco, T. M. Jessell, K. A. Wilkinson, and A. Patapoutian, 2015. Piezo2 is the principal mechanotransduction channel for proprioception. Nat. Neurosci. 18:1756–1762.

16. Szczot, M., J. Liljencrantz, N. Ghitani, A. Barik, R. Lam, J. H. Thompson, D. Bharucha-Goebel, D. Saade, A. Necaise, S. Donkervoort, A. R. Foley, T. Gordon, L. Case, M. C. Bushnell, C. G. Bönnemann, and A. T. Chesler, 2018. PIEZO2 mediates injury-induced tactile pain in mice and humans. Sci. Transl. Med. 10:eaat9892.

17. Murthy, S. E., M. C. Loud, I. Daou, K. L. Marshall, F. Schwaller, J. Kühnemund, A. G. Francisco, W. T. Keenan, A. E. Dubin, G. R. Lewin, and A. Patapoutian, 2018. The mechanosensitive ion channel Piezo2 mediates sensitivity to mechanical pain in mice. Sci. Transl. Med. 10:aat9897.

18. Holt, J. R., W.-Z. Zeng, E. L. Evans, S.-H. Woo, S. Ma, H. Abuwarda, M. Loud, A. Patapoutian, and M. M. Pathak, 2021. Spatiotemporal dynamics of PIEZO1 localization controls keratinocyte migration during wound healing. eLife 10:e65415.

19. Pathak, M. M., J. L. Nourse, T. Tran, J. Hwe, J. Arulmoli, D. T. T. Le, E. Bernardis, L. A. Flanagan, and F. Tombola, 2014. Stretch-activated ion channel Piezo1 directs lineage choice in human neural stem cells. Proc. Natl. Acad. Sci. U.S.A. 111:16148–16153.

20. Jankovsky, N., A. Caulier, J. Demagny, C. Guitton, S. Djordjevic, D. Lebon, H. Ouled-Haddou, V. Picard, and L. Garçon, 2021. Recent advances in the pathophysiology of PIEZO1-related hereditary xerocytosis. Am. J. Hematol. 96:1017–1026.

21. Lukacs, V., J. Mathur, R. Mao, P. Bayrak-Toydemir, M. Procter, S. M. Cahalan, H. J. Kim, M. Bandell, N. Longo, R. W. Day, D. A. Stevenson, A. Patapoutian, and B. L. Krock, 2015. Impaired PIEZO1 function in patients with a novel autosomal recessive congenital lymphatic dysplasia. Nat. Comm. 6:6:8329.

22. Saotome, K., S. E. Murthy, J. M. Kefauver, T. Whitwam, A. Patapoutian, and A. B. Ward, 2017. Structure of the mechanically activated ion channel Piezo1. Nature 554:481–486.

23. Ge, J., W. Li, Z. Qiancheng, N. Li, M. Chen, P. Zhi, R. Li, N. Gao, B. Xiao, and M. Yang, 2015. Architecture of the mammalian mechanosensitive Piezo1 channel. Nature 527:64–69.

24. Guo, Y. R., and R. MacKinnon, 2017. Structure-based membrane dome mechanism for Piezo mechanosensitivity. eLife 6:e33660.

25. Zhao, Q., H. Zhou, S. Chi, Y. Wang, J. Wang, J. Geng, K. Wu, W. Liu, T. Zhang, M.-Q. Dong, J. Wang, X. Li, and B. Xiao, 2018. Structure and mechanogating mechanism of the Piezo1 channel. Nature 554:487–492.

26. Wang, L., H. Zhou, M. Zhang, W. Liu, T. Deng, Q. Zhao, Y. Li, J. Lei, X. Li, and B. Xiao, 2019. Structure and mechanogating of the mammalian tactile channel PIEZO2. Nature 573:225–229. 10.1038/s41586-019-1505-8.

27. Haselwandter, C. A., and R. MacKinnon, 2018. Piezo’s membrane footprint and its contribution to mechanosensitivity. eLife 7:e41968.

28. Mulhall, E. M., A. Gharpure, R. M. Lee, A. E. Dubin, J. S. Aaron, K. L. Marshall, K. R. Spencer, M. A. Reiche, S. C. Henderson, T.-L. Chew, and A. Patapoutian, 2023. Direct observation of the conformational states of PIEZO1. Nature 620:1117–1125. 10.1038/s41586-023-06427-4.

29. Lin, Y.-C., Y. R. Guo, A. Miyagi, J. Levring, R. MacKinnon, and S. Scheuring, 2019. Force-induced conformational changes in PIEZO1. Nature 573:230–234. 10.1038/s41586-019-1499-2.

30. Vero Li, J., C. D Cox, and B. Martinac, 2021. The anchor domain is critical for Piezo1 channel mechanosensitivity. Channels 15:438–446. 10.1080/19336950.2021.1923199.

31. Lewis, A. H., and J. Grandl, 2015. Mechanical sensitivity of Piezo1 ion channels can be tuned by cellular membrane tension. eLife 4:e12088.

32. Syeda, R., M. N. Florendo, C. D. Cox, J. M. Kefauver, J. S. Santos, B. Martinac, and A. Patapoutian, 2016. Piezo1 Channels Are Inherently Mechanosensitive. Cell Rep. 17:1739–1746.

33. Cox, C. D., C. Bae, L. Ziegler, S. Hartley, V. Nikolova-Krstevski, P. R. Rohde, C.-A. Ng, F. Sachs, P. A. Gottlieb, and B. Martinac, 2016. Removal of the mechanoprotective influence of the cytoskeleton reveals PIEZO1 is gated by bilayer tension. Nat. Commun. 7:10366.

34. Bavi, N., J. Richardson, C. Heu, B. Martinac, and K. Poole, 2019. PIEZO1-Mediated Currents Are Modulated by Substrate Mechanics. ACS Nano 13:13545–13559.

35. Wang, J., J. Jiang, X. Yang, G. Zhou, L. Wang, and B. Xiao, 2022. Tethering Piezo channels to the actin cytoskeleton for mechanogating via the cadherin-*β*-catenin mechanotransduction complex. Cell Rep. 38:110342.

36. Ellefsen, K. L., J. R. Holt, A. C. Chang, J. L. Nourse, J. Arulmoli, A. H. Mekhdjian, H. Abuwarda, F. Tombola, L. A. Flanagan, A. R. Dunn, I. Parker, and M. M. Pathak, 2019. Myosin-II mediated traction forces evoke localized Piezo1-dependent Ca2 flickers. Commun. Biol. 2:298.

37. Ridone, P., E. Pandzic, M. Vassalli, C. D. Cox, A. Macmillan, P. A. Gottlieb, and B. Martinac, 2020. Disruption of membrane cholesterol organization impairs the activity of PIEZO1 channel clusters. J. Gen. Physiol. 152:e201912515.

38. Sezgin, E., I. Levental, S. Mayor, and C. Eggeling, 2017. The mystery of membrane organization: composition, regulation and roles of lipid rafts. Nat. Rev. Mol. Cell Biol. 18:361–374.

39. Jacobson, K., P. Liu, and B. C. Lagerholm, 2019. The Lateral Organization and Mobility of Plasma Membrane Components. Cell 177:806–819.

40. Singer, S. J., and G. L. Nicolson, 1972. The Fluid Mosaic Model of the Structure of Cell Membranes: Cell membranes are viewed as two-dimensional solutions of oriented globular proteins and lipids. Science 175:720–731. 10.1126/science.175.4023.720.

41. Vaisey, G., P. Banerjee, A. J. North, C. A. Haselwandter, and R. MacKinnon, 2022. Piezo1 as a force-through-membrane sensor in red blood cells. eLife 11. 10.7554/elife.82621.

42. Edelstein, A., N. Amodaj, K. Hoover, R. Vale, and N. Stuurman, 2010. Computer Control of Microscopes Using μManager. Curr. Protoc. Mol. Biol. 92:14.20.1––14.20.17.

43. Ellefsen, K. L., B. Settle, I. Parker, and I. F. Smith, 2014. An algorithm for automated detection, localization and measurement of local calcium signals from camera-based imaging. Cell Calcium 56:147–156.

44. Harris, C. R., K. J. Millman, S. J. van der Walt, R. Gommers, P. Virtanen, D. Cournapeau, E. Wieser, J. Taylor, S. Berg, N. J. Smith, R. Kern, M. Picus, S. Hoyer, M. H. van Kerkwijk, M. Brett, A. Haldane, J. F. del Río, M. Wiebe, P. Peterson, P. Gérard-Marchant, K. Sheppard, T. Reddy, W. Weckesser, H. Abbasi, C. Gohlke, and T. E. Oliphant, 2020. Array programming with NumPy. Nature 585:357–362.

45. R Core Team, 2016. R: A Language and Environment for Statistical Computing. R Foundation for Statistical Computing, Vienna, Austria. https://www.R-project.org/.

46. Golan, Y., and E. Sherman, 2017. Resolving mixed mechanisms of protein subdiffusion at the T cell plasma membrane. Nat. Commun. 8:1–15.

47. Benaglia, T., D. Chauveau, D. R. Hunter, and D. Young, 2009. mixtools: An R Package for Analyzing Finite Mixture Models. Journal of Statistical Software 32:1–29. https://www.jstatsoft.org/v32/i06/.

48. Inc., W. R. Mathematica, Version 14.0. Champaign, IL, 2024.

49. Kepten, E., I. Bronshtein, and Y. Garini, 2013. Improved estimation of anomalous diffusion exponents in single-particle tracking experiments. Phys. Rev. E 87:052713.

50. Martin-Gonthier, P., E. Havard, and P. Magnan, 2010. Custom transistor layout design techniques for random telegraph signal noise reduction in CMOS image sensors. Electronics Letters 46:1323. 10.1049/el.2010.1767.

51. Janesick, J. R., T. Elliott, S. Collins, M. M. Blouke, and J. Freeman, 2001. Scientific Charge-Coupled Devices. Optical Engineering 26. 10.1117/12.7974139.

52. Huang, F., T. M. P. Hartwich, F. E. Rivera-Molina, Y. Lin, W. C. Duim, J. J. Long, P. D. Uchil, J. R. Myers, M. A. Baird, W. Mothes, M. W. Davidson, D. Toomre, and J. Bewersdorf, 2013. Video-rate nanoscopy using sCMOS camera–specific single-molecule localization algorithms. Nature Methods 10:653–658. 10.1038/nmeth.2488.

53. Yao, M., A. Tijore, D. Cheng, J. V. Li, A. Hariharan, B. Martinac, G. T. V. Nhieu, C. D. Cox, and M. Sheetz, 2022. Force- and cell state–dependent recruitment of Piezo1 drives focal adhesion dynamics and calcium entry. Science Advances 8. 10.1126/sciadv.abo1461.

54. Gudipaty, S. A., J. Lindblom, P. D. Loftus, M. J. Redd, K. Edes, C. F. Davey, V. Krishnegowda, and J. Rosenblatt, 2018. A Feedforward Mechanism Mediated by Mechanosensitive Ion Channel PIEZO1 and Tissue Mechanics Promotes Glioma Aggression. Neuron 100:799–815.e7.

55. He, W., H. Song, Y. Su, L. Geng, B. J. Ackerson, H. B. Peng, and P. Tong, 2016. Dynamic heterogeneity and non-Gaussian statistics for acetylcholine receptors on live cell membrane. Nat. Comm. 7.

56. Rehfeldt, F., and M. Weiss, 2023. The random walker’s toolbox for analyzing single-particle tracking data. Soft Matter 19:5206–5222. 10.1039/d3sm00557g.

57. Akaike, H., 1974. A new look at the statistical model identification. IEEE Transactions on Automatic Control 19:716–723. 10.1109/TAC.1974.1100705.

58. Nicolson, G. L., 2014. The Fluid—Mosaic Model of Membrane Structure: Still relevant to understanding the structure, function and dynamics of biological membranes after more than 40years. Biochim. Biophys. Acta 1838:1451–1466.

59. Bohórquez-Hernández, A., E. Gratton, J. Pacheco, A. Asanov, and L. Vaca, 2017. Cholesterol modulates the cellular localization of Orai1 channels and its disposition among membrane domains. Biochim. Biophys. Acta 1862:1481–1490.

60. Abu-Arish, A., E. Pandzic, J. Goepp, E. Matthes, J. W. Hanrahan, and P. W. Wiseman, 2015. Cholesterol Modulates CFTR Confinement in the Plasma Membrane of Primary Epithelial Cells. Biophys. J. 109:85–94.

61. Adkins, E. M., D. J. Samuvel, J. U. Fog, J. Eriksen, L. D. Jayanthi, C. B. Vaegter, S. Ramamoorthy, and U. Gether, 2007. Membrane Mobility and Microdomain Association of the Dopamine Transporter Studied with Fluorescence Correlation Spectroscopy and Fluorescence Recovery after Photobleaching. ACS Biochemistry 46:10484–10497.

62. Romero, L. O., A. E. Massey, A. D. Mata-Daboin, F. J. Sierra-Valdez, S. C. Chauhan, J. F. Cordero-Morales, and V. Vásquez, 2019. Dietary fatty acids fine-tune Piezo1 mechanical response. Nat. Commun. 10:1200.

63. Bae, C., F. Sachs, and P. A. Gottlieb, 2011. The mechanosensitive ion channel Piezo1 is inhibited by the peptide GsMTx4. Biochemistry 50:6295–6300.

64. Höfling, F., and T. Franosch, 2013. Anomalous transport in the crowded world of biological cells. Rep. Prog. Phys. 76:046602.

65. Metzler, R., J.-H. Jeon, and A. Cherstvy, 2016. Non-Brownian diffusion in lipid membranes: Experiments and simulations. Biochim. Biophys. Acta 1858:2451–2467.

66. Martin, D. S., M. B. Forstner, and J. A. Käs, 2002. Apparent Subdiffusion Inherent to Single Particle Tracking. Biophys. J. 83:2109–2117.

67. Mainali, D., A. Syed, N. Arora, and E. A. Smith, 2014. Role of insulin receptor and insulin signaling on *α*PS2C*β*PS integrins’ lateral diffusion. European Biophysics Journal 43:603–611. 10.1007/s00249-014-0990-9.

68. Srinivasan, Y., A. Guzikowski, R. Haugland, and K. Angelides, 1990. Distribution and lateral mobility of glycine receptors on cultured spinal cord neurons. The Journal of Neuroscience 10:985–995. 10.1523/JNEUROSCI.10-03-00985.1990.

69. Sanfeliu-Cerdán, N., F. Català-Castro, B. Mateos, C. Garcia-Cabau, M. Ribera, I. Ruider, M. Porta-de-la Riva, A. Canals-Calderón, S. Wieser, X. Salvatella, and M. Krieg, 2023. A MEC-2/stomatin condensate liquid-to-solid phase transition controls neuronal mechanotransduction during touch sensing. Nature Cell Biology 25:1590–1599. 10.1038/s41556-023-01247-0.

70. Nicoll, R. A., 2017. A Brief History of Long-Term Potentiation. Neuron 93:281–290. 10.1016/j.neuron.2016.12.015.

71. Bailey, D. M., M. A. Catron, O. Kovtun, R. L. Macdonald, Q. Zhang, and S. J. Rosenthal, 2018. Single Quantum Dot Tracking Reveals Serotonin Transporter Diffusion Dynamics are Correlated with Cholesterol-Sensitive Threonine 276 Phosphorylation Status in Primary Midbrain Neurons. ACS Chem. Neurosci 9:2534–2541.

72. Poole, K., R. Herget, L. Lapatsina, H.-D. Ngo, and G. R. Lewin, 2014. Tuning Piezo ion channels to detect molecular-scale movements relevant for fine touch. Nature Communications 5. 10.1038/ncomms4520.

73. Qi, Y., L. Andolfi, F. Frattini, F. Mayer, M. Lazzarino, and J. Hu, 2015. Membrane stiffening by STOML3 facilitates mechanosensation in sensory neurons. Nat. Comm. 6. 10.1038/ncomms9512.

74. Mott, T. M., G. C. Wulffraat, A. J. Eddins, R. A. Mehl, and E. N. Senning, 2024. Fluorescence labeling strategies for cell surface expression of TRPV1. Journal of General Physiology 156. 10.1085/jgp.202313523.

75. Varoquaux, G., L. Buitinck, G. Louppe, O. Grisel, F. Pedregosa, and A. Mueller, 2015. Scikit-learn: Machine Learning Without Learning the Machinery. GetMobile: Mobile Computing and Communications 19:29–33. 10.1145/2786984.2786995.

